# Understanding potential drivers of aquatic metabolism in a subtropical treatment wetland

**DOI:** 10.1101/2020.05.02.074310

**Authors:** Paul Julian, Todd Z. Osborne

## Abstract

Changes of dissolved oxygen (DO) in aquatic ecosystems integrates dynamic biological, physical and chemical processes that control the rate of ecosystem metabolism. Aquatic ecosystem metabolism can be characterized by the diel change in DO changes over time and is expressed as the net aquatic productivity (NAP). This study investigated aquatic metabolism of dominant emergent and submerged aquatic vegetation (EAV and SAV, respectively) within two treatment flow-ways (FW) of Stormwater Treatment Area 2 (STA-2) in the Everglades ecosystem. The hypothesis of this study is that aquatic metabolism will differ between aquatic vegetation communities with SAV communities will have a greater GPP and ER rate than EAV communities driven by biophysical, hydrodynamic and biogeochemical differences between systems. Aquatic metabolism observed in this study vary spatially (along FWs) and temporally (diel to days) controlled by different effects related biological, physical and chemical processes. This study suggests that ecosystem metabolism is controlled differently across FWs with varying levels of response to loading/transport and water column attributes resulting in differences in organic matter accumulation, C turnover and phosphorus cycling.

## Introduction

> *“To the casual eye the biota of flowing waters is rich.”*
>
> — - (Odum 1956)

The measurement of production and oxidation of organic matter forms the basic understanding of ecosystem function. In lotic ecosystems as part of the river continuum concept Vannote et al. (1980) hypothesized that gradual shifts in the ratio of gross primary productivity (GPP) to ecosystem respiration (ER) occur as rivers transition from headwaters to mid-sized streams due to changes to ecosystem geomorphology along the river and processing of detrital material by ecosystem components (i.e. benthic macroinvertebrates). Generally in lotic ecosystem (i.e. streams and rivers) metabolic behavior (i.e. GPP and ER) is driven largely by hydrodynamic variability (i.e. load) resulting in relatively slow mineralization rates (Hotchkiss et al. 2018). Meanwhile, in lentic ecosystems (lakes) metabolic behavior is regulated by both autochthonous and allochthonous carbon (C) and growth limiting nutrients such as phosphorus (P) (del Giorgio and Peters 1994; Carignan et al. 2000; Cole et al. 2000; Hanson et al. 2003). In wetland ecosystems, which straddle the lentic-lotic divide, in general metabolically resemble lake ecosystems where both auto- and allochthonous C drive ecosystem metabolism (McKenna 2003; Kadlec and Wallace 2009; Hagerthey et al. 2010) with some evidence suggesting that hydrologic variability can be also contribute to changes in metabolism (Tuttle et al. 2008; Hornbach et al. 2017).

Wetlands are generally net C sinks that store a large amount of the global C, created through an unbalanced accumulation of C through plant productivity and export from decomposition of organic matter via gaseous C (i.e. C dioxide and methane). Dissolved and particulate C in the water column can be transported laterally through run-off providing allochthonous subsidies of C to downstream systems (Updegraff et al. 1995; Carpenter and Pace 1997; Freeman et al. 2004; Billett and Moore 2008). Hydrologic conditions, nutrient input and nutrient cycling regulate the C balance, speciation and flux C from wetland ecosystems. Nutrient inputs and hydrologic conditions regulate the balance between dissolve inorganic and dissolved organic C (DIC and DOC, respectively) production via net productivity and metabolism of organic matter (Julian et al. 2017). Generally, C dynamics within treatment wetlands are similar to that of natural wetlands however, treatment wetlands typically receive proportionally larger allochthonous C inputs depending on landscape position and upstream watershed characteristics (Reddy and DeLaune 2008; Kadlec and Wallace 2009). In treatment wetlands used to treat wastewater effluent, the sequestration of C and the associated reduction in biological oxygen demand can be substantial with some seasonal variability controlled largely by climate, plant biomass cycling and water temperatures (Kadlec and Wallace 2009).

Treatment wetlands are tools to provide natural removal of water column pollutants and ultimately improve water quality conditions for downstream ecosystems. In the south Florida, treatment wetlands were constructed to provide water quality improvements and restore the biological integrity of the Everglades ecosystem. These treatment wetlands were constructed to improve the quality of agricultural runoff water originating in the Everglades Agricultural Area (EAA) prior to entering the downstream Everglades ecosystem (Chen et al. 2015). These treatment wetlands, referred to as the Everglades stormwater treatment areas (STAs) were constructed with the primary objective of removing excess P from surface water prior to discharge to the Everglades Protection Area. The Everglades ecosystem is a P-growth limited system with flora and fauna adapted to nutrient poor (i.e. oligotrophic) conditions (Noe et al. 2001). The STAs are composed of several treatment cells or flow-ways (FWs) which use natural wetland communities to facilitate the removal of P from the water column by leveraging natural wetland processes including nutrient storage into vegetative tissues and burial within soils (Kadlec and Wallace 2009). While the focus of the Everglades STAs is P, it is possible that P, nitrogen (N) and C cycles are coupled sharing co-dependencies and interrelationships (Corstanje et al. 2016; Julian et al. 2019).

Given these possible co-dependencies and inter-relationships between P and C, understanding these relationships could gain a deeper understanding of system performance and outcomes. The objects of this study were to 1) evaluate aquatic metabolism within and between FWs with contrasting dominate aquatic vegetation communities specifically to consider soil C dynamics and 2) explore biotic and abiotic drivers of aquatic GPP in a shallow, subtropical treatment wetland. The first hypothesis is that aquatic metabolism will differ between aquatic vegetation communities with submerged aquatic vegetation (SAV) communities will have a greater GPP and ER rate than emergent aquatic vegetation (EAV) communities. The second hypothesis is that hydrologic loading and water column characteristics such as Total P (TP), Chlorophyll *a,* DOC and turbidity will regulate GPP to some degree in both FWs with nutrients and algal biomass providing up-regulation of GPP and optical water quality parameters (turbidity, chlorophyll-*a*, etc.) down regulating GPP.

## Material and methods

### Study Area

A total of six STAs with an approximate area of 231 km^2^ are located south of Lake Okeechobee in the southern portion of the Everglades Agricultural Area (Fig. 1). The Everglades STAs are established on land which earlier had natural wetlands and/or agricultural farmlands under sugarcane production. The primary source of inflow water to the STAs is agricultural runoff originating from approximately 284 km^2^ of farmland upstream. Everglades STA treatment cells or FWs are comprised of a mixture of EAV and SAV communities in several configurations including EAV and SAV treatment cells arranged in parallel or in series (Chen et al. 2015).

**Figure 1.**
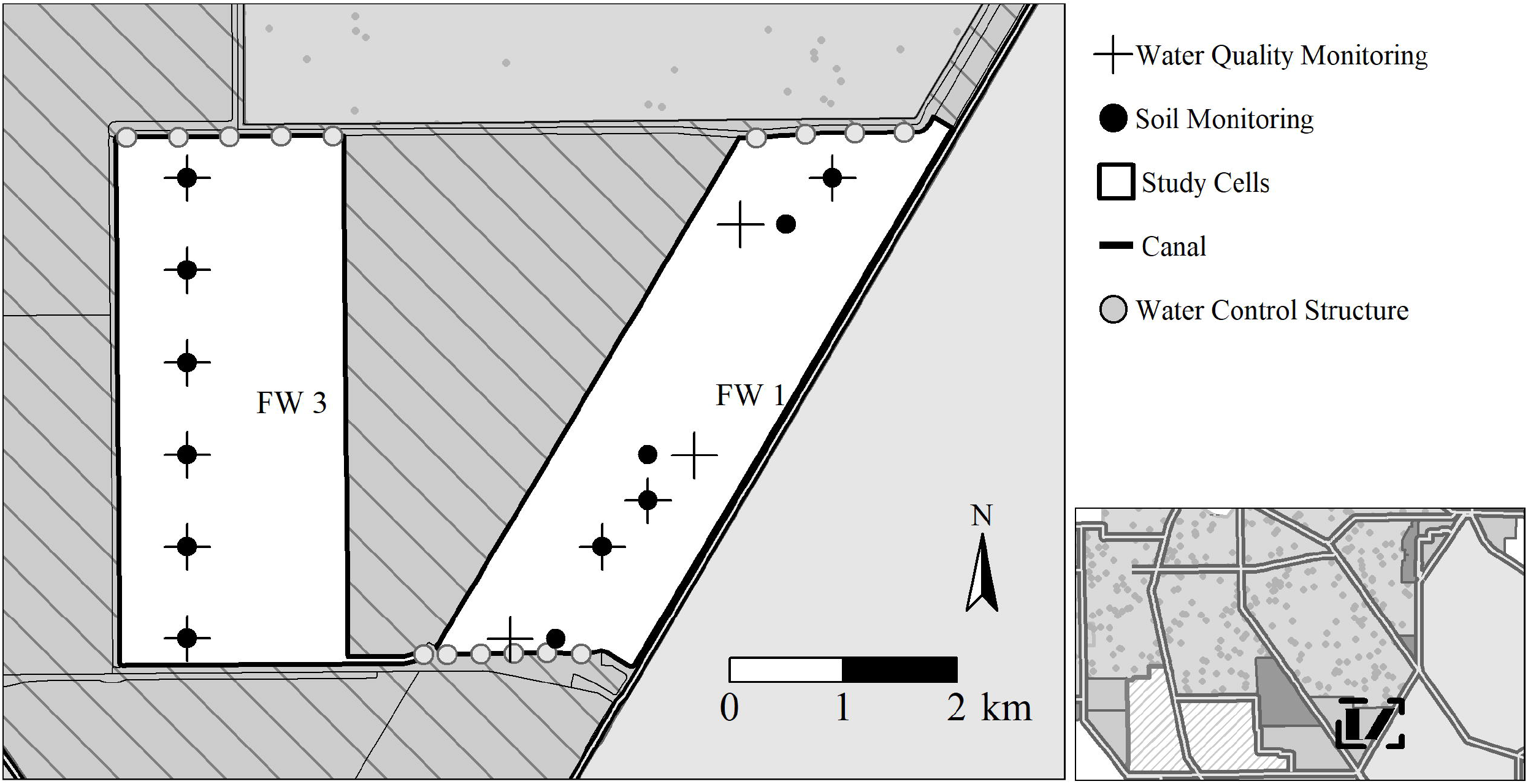
Surface water, soil and vegetation monitoring locations within Everglades Stormwater Treatment Area-2 Cells 1 (right) and 3 (left). Cell 1 is predominately emergent vegetation and Cell 3 is predominately submerged aquatic vegetation. Operationally these cells are identified as flow-way 1 and 3, respectively.

Stormwater Treatment Area-2 has been in operation since June 1999 with an effective treatment area of approximately 63 km^2^ divided into eight treatment cells. This study was conducted in two cells, FWs 1 and 3. The vegetative community of FW 1 is comprised predominately of EAV vegetation including *Typha domingensis* Pers. (cattail) and *Cladium jamaicense* Crantz (sawgrass) while FW 3 is dominantly SAV including *Chara* spp. (muskgrass), *Potamogeton* spp. (pondweed) and *Najas guadalupensis* Spreng (southern naiad) with approximately a third of the FW occupied by EAV species. Furthermore, prior to STA-2 construction FW 1 was a historic natural wetland while approximate two-thirds of FW 3 was previously farmed and is now managed as a SAV system (Juston and DeBusk 2006).

### Data Source

Data used in this study were collected by South Florida Water Management District (SFWMD) and University of Florida as a part of a larger project within the overall SFWMD’s Restoration Strategies Science Plan to evaluate performance and optimization of the Everglades STAs (South Florida Water Management District 2013). Data from one of the main studies of the Science Plan to evaluate P-sources, form, fluxes and transformation process in the STAs was used for this investigation (UF-WBL 2017; Villapando and King 2018). Water quality monitoring locations were established along two FWs within STA-2 along a transect running from inflow to outflow of the FW (Fig 1). Water quality sondes were deployed at established monitoring locations and programed to collect dissolved oxygen (DO), temperature, pH and specific conductivity and turbidity data every 15 to 30 minutes depending on deployment duration. Data from deployed sondes underwent a two-tier quality assurance/quality control (QA/QC) consistent with method used by the National Estuary Research Reserve Program (NERR) following the data management protocols developed by the Centralized Data Management Office (NOAA 2013).

Weekly surface water grab samples collected at monitoring locations within FWs 1 and 3 were used to characterize changes in nutrient concentrations during prescribed/semi-managed flow event. Flow events were planned as a short duration (fixed temporal window) events which flows to the system were maintained within a pre-determined range, and extensive monitoring was undertaken to ascertain system’s response to the controlled flow regime. Water quality samples collected along a gradient from inflow to outflow were analyzed for TP, chlorophyll *a*, and DOC (Table S1). Flocculent and soil samples were collected along the flow transects twice during the dry and wet seasons between 2015 and 2016. Soils were sampled using the push-core method consistent with methods used for prior wetland soil studies (Bruland et al. 2007; Osborne et al. 2011; Newman et al. 2017). A 10-cm diameter polycarbonate core tube was pushed into the soil until refusal. The consolidated soil was segmented with the 0 - 5 cm segment retained for analyses. Soil samples were analyzed for several parameters but for purposes of this study only bulk density, total C (TC) and loss-on-ignition (LOI) were considered (Table S1). High resolution weather data were retrieved from the SFWMD online environmental database (DBHYDRO; www.sfwmd.gov/dbhydro) from two weather stations (STA1W and ROTNWX) located within close proximity of the study area (Fig 1). Surface flow volume and total TP concentrations at the inflow and outflow of each FW were also retrieved from DBHYDRO during each flow event.

### Data Analysis

Aquatic metabolism was estimated using hourly mean DO, surface water temperature, wind speed, air temperature and barometric pressure. Gross primary productivity (GPP), ecosystem respiration (ER) and net aquatic productivity (NAP) were estimated using the “open-water method” first proposed by Odum (1956) and employed by others (Cole et al. 2000; Thébault et al. 2008; Hagerthey et al. 2010; Staehr et al. 2010; Caffrey et al. 2014). Daily NAP, GPP and ER were calculated on the basis of changes in DO concentrations assumed to be driven by rate of photosynthesis, respiration and atmospheric exchange (Eq 1; Odum 1956). Net aquatic productivity and ER can be determined directly from DO sensor data, GPP is estimated by mass-balance. Hourly mean DO concentrations were calculated from the 15 – 30 minute data to provide a basis to compare changes in DO over consistent time-steps. Within a 1-h interval, it is assumed that the change in DO is equal to the sum of the respiration rate and oxygen diffusion rate minus the rate of photosynthesis (Eq. 1) where C is the dissolved oxygen concentration (mg L^−1^), t is time (h), P is the rate of photosynthesis (mg L^−1^ h^−1^), R is the respiration rate (mg L^−1^ h^−1^), and D is the rate of oxygen uptake from diffusion across the air-water interface (mg L^−1^ h^−1^).

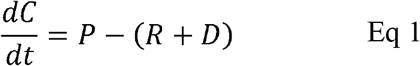

The rate of oxygen uptake by diffusion across the air-water interface (D) is regulated by the difference in O_2_ in the water column from atmospheric equilibrium and the temperature-dependent gas exchange coefficient for oxygen. Wind produces turbulence in stationary water bodies, facilitating gas exchange processes driven by wind speed (Eq. 2) where k_a_ is the volumetric re-aeration coefficient (h^−1^), C_s_ is the DO saturation concentration (mg L^−1^) and C is the dissolved oxygen concentration (mg L^−1^). It is assumed that the wind speed measured at the nearby meteorological stations would be a reasonable surrogate for the wind conditions for sampling stations location in open water (i.e. FW 3) areas but not for sites with dense emergent vegetation (i.e. FW 1). For FW 1, it was assumed that the emergent vegetation would reduce the wind speed the air-water interrace to effectively zero consistent with Hagerthey et al. (2010). The volumetric re-aeration coefficient was calculated using three different functions of wind speed as an approach for estimating uncertainty in the rate of air-water oxygen exchange consistent with prior studies (Thébault et al. 2008; Caffrey et al. 2014).

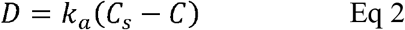

Net aquatic productivity is the sum of photosynthesis and respiration which can be computed by Eq. 3. For each day, NAP was calculated by summing hourly diffusion-corrected rate of DO change over a 24-h period, starting and ending at sunrise. Sunrise and sunset was based on monitoring location longitude and latitude and estimated using algorithms developed by NOAA in a modified version of the “metab_day” function in the ‘SWMPr’ R-package (Beck 2016) and verified with light intensity data retrieved from near-by weather stations. Hourly ER was computed as the mean night-time diffusion corrected rates of DO changes, it was assumed that GPP was zero at night (Cole et al. 2000) and the change in DO within an hourly interval during the night is attributed to respiration and diffusion. Assuming that nighttime ER is equal to daytime ER, daily respiration was calculated by multiplying hourly ER by 24 h. Hourly GPP was calculated by subtracting ER from diffusion corrected rates of DO change during the daylight period while daily photosynthesis was computed by summing hourly photosynthesis values from sunrise to sunset. Volumetric rates of GPP, ER, and NAP were multiplied by depth to yield areal productivity rates (g O_2_ m^−2^ d^−1^) consistent with methods employed by other studies (Cole et al. 2000; Thébault et al. 2008; Hagerthey et al. 2010; Staehr et al. 2010; Caffrey et al. 2014). Metabolism calculations were performed using a modified version of the “metabolism” function in the ‘SWMPr’ R-package (Beck 2016).

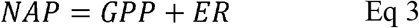

Areal soil C storage expressed as g C m^−2^ was calculated following Eq 4 where TC is total carbon concentration (mg kg^−1^), BD is bulk density (g cm^−3^) and d is the soil layer depth (cm).

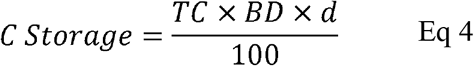

Hydraulic loading rate (HLR) was calculated for each flow event based on methods by Kadlec and Wallace (2009). Daily HLR values were derived for each FW by dividing daily total inflow volume and FW area.

Chi-squared (R x C) analysis was used to compare hourly averaged DO concentrations by FW to evaluate overall diel changes in DO. Daily GPP and ER values were compared between transect location for each FW using the Kruskal-Wallis rank sum test and Dunn’s test of multiple comparison (Dinno 2015). Spearman’s rank correlation test was used to compare daily spatially averaged GPP, ER and NAP for each FW versus FW specific daily HLR during each flow event. The ratio between GPP and ER was compared to one for each FW using one-sample Wilcoxon signed rank test. Spearman’s rank correlation test was also used to GPP and water quality parameters on days water quality grab samples were collected for each FW. Water quality parameters used in the comparison include TP, chlorophyll *a* and daily mean turbidity (from sonde). Due to sample size limitations with respect to soil C, qualitative comparison of aquatic metabolism (i.e. GPP, ER and NAP) relative to soil C were conducted.

Unless otherwise noted, all statistical operations were performed using the base stats R-package. All statistical operations were performed using R (Ver 3.5.2, R Foundation for Statistical Computing, Vienna Austria) and the critical level of significance was set at α = 0.05.

## Results and discussion

Natural wetlands generally have a high rate of biological activity, more than most other ecosystems. Due to this high biological activity, natural wetland processes can be leveraged to facilitate the removal of contaminants in man-made constructed treatment wetland systems. During these processes, contaminants are either transformed to harmless byproducts or essential nutrients that can be used for additional biological productivity (Kadlec and Wallace 2009). The Everglades stormwater treatment areas were constructed to remove excess P from agricultural runoff prior to discharging that water to the downstream Everglades ecosystem (Chen et al. 2015). Numerous studies have explored and evaluated the P-removal capacity of these systems to protect the Everglades ecosystem from large-scale nutrient enrichment and degradation (Chimney and Goforth 2001; Newman and Pietro 2001; DeBusk et al. 2004; Juston and DeBusk 2011; Reddy et al. 2011; Corstanje et al. 2016) however, little attention has been given to the role aquatic metabolism relative to these processes.

Using high temporal resolution DO, temperature and weather data, estimates of aquatic metabolism were determined for a total of 330 days at 15 sites across two STA FWs during six semi-prescribed flow events during this study. Although the equipment was deployed during the short duration flow events, various issues (i.e. biofouling, wildlife interactions, sensor failure, etc.) produced data gaps throughout our record. Across all sites, we were able to estimate GPP and ER for 79% of the total possible sampling days.

### Diurnal DO Responses

Across the two FWs during the total of six flow events, hourly mean DO concentrations range from anoxic (0 mg O_2_ L^−1^) to over 38 mg O_2_ L^−1^. A strong diurnal response in DO concentrations was observed across all sites and FWs with minimum DO concentrations occurring during the early morning hours, beginning to climb around mid-day and peaking after sunset (Fig 2). Hourly DO concentrations were significantly greater in FW 3 (χ^2^ = 14416, df=1, ρ<0.01) when compared to FW 1 (Fig 2). The dominate vegetative community of FW 3 SAV contribute significantly to the oxygenation of water column and aquatic GPP more so than EAV (Caraco and Cole 2002; McCormick and Laing 2003; Hagerthey et al. 2010). As its names implies, SAV is submerged in the water column therefore any gas exchange by the plants is done predominately in the water column, as opposed to EAV where only a portion of the plant is submerged. Water column oxygenation by SAV is facilitated by photosynthetically-produced oxygen diffusing through the plant and can be enhanced by light availability as observed in lake, stream and wetland ecosystems (Caraco and Cole 2002; Caraco et al. 2006; Hagerthey et al. 2010). As such, this exchange of oxygen into the ambient environment can significantly influence the oxidation-reduction (redox) conditions near the vegetation particularly in the soil around the rhizomes supplementing sediment oxygen demand needs(Reddy and DeLaune 2008). In addition to changes in water column oxygen, concurrent changes in water column pH are notable in SAV communities (Fig 3 and Table 1) where the photosynthesis and respiration by SAV strongly influences pH and inorganic carbon speciation and flux (Carpenter and Lodge 1986; Sand-Jensen and Frost-Christensen 1998; Julian et al. 2017). These changes in DO, redox conditions and the pH environment can affect P sources and sinks through the precipitation of minerals including phosphate and calcium minerals (Sand-Jensen et al. 1982; Carpenter and Lodge 1986; Harris 2011) and chemical oxidation of organic matter (Liao and Gurol 1995).

**Figure 2.**
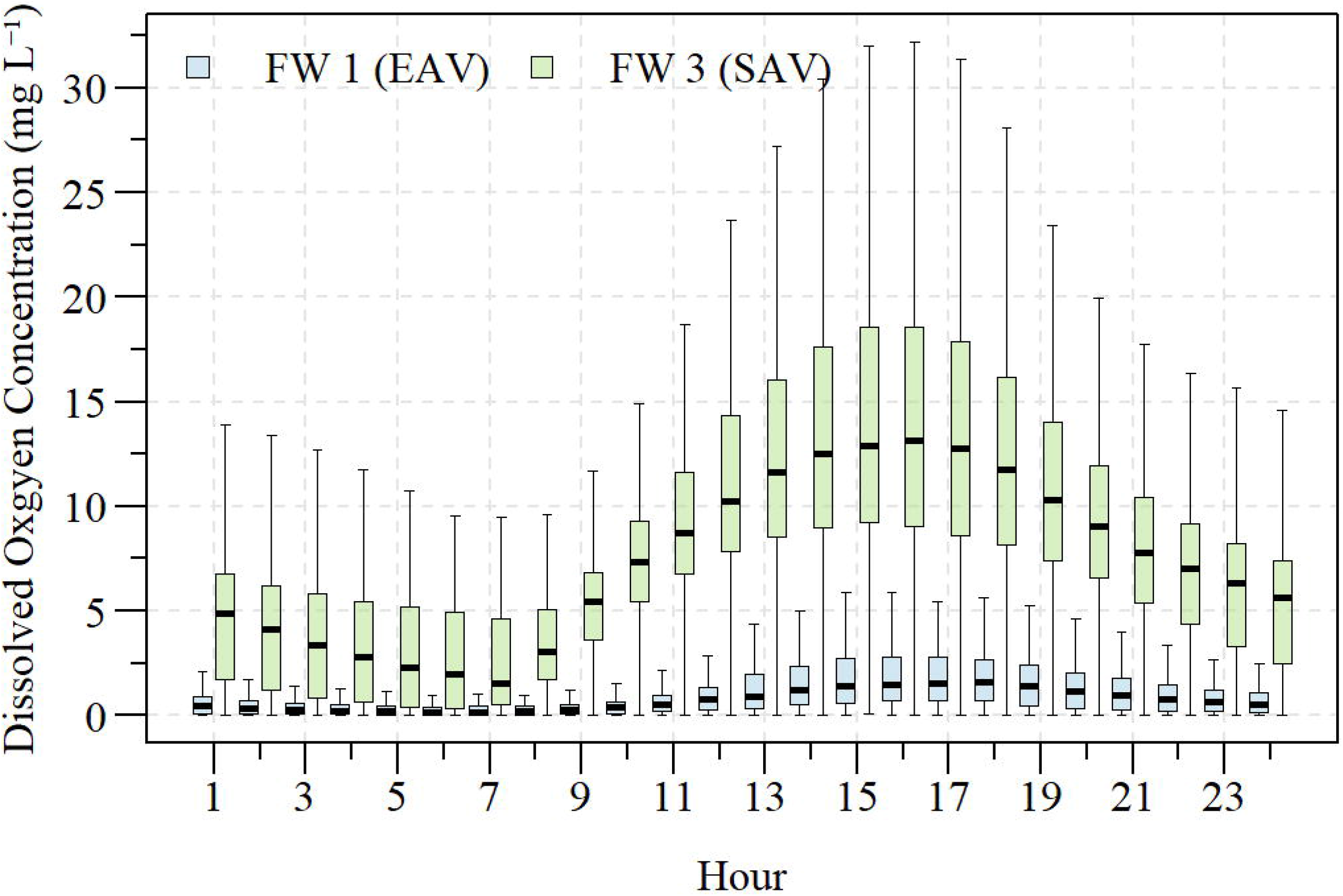
Diurnal dissolved oxygen concentration for all sites along the flow-way (FW) 1 and FW 3 transects within Stormwater Treatment Area 2 measured during this study. Hour represented in 24-hour clock. Dominate vegetation of FW 1 and 3 is emergent aquatic vegetation (EAV) and submerged aquatic vegetation (SAV), respectively.

**Figure 3.**
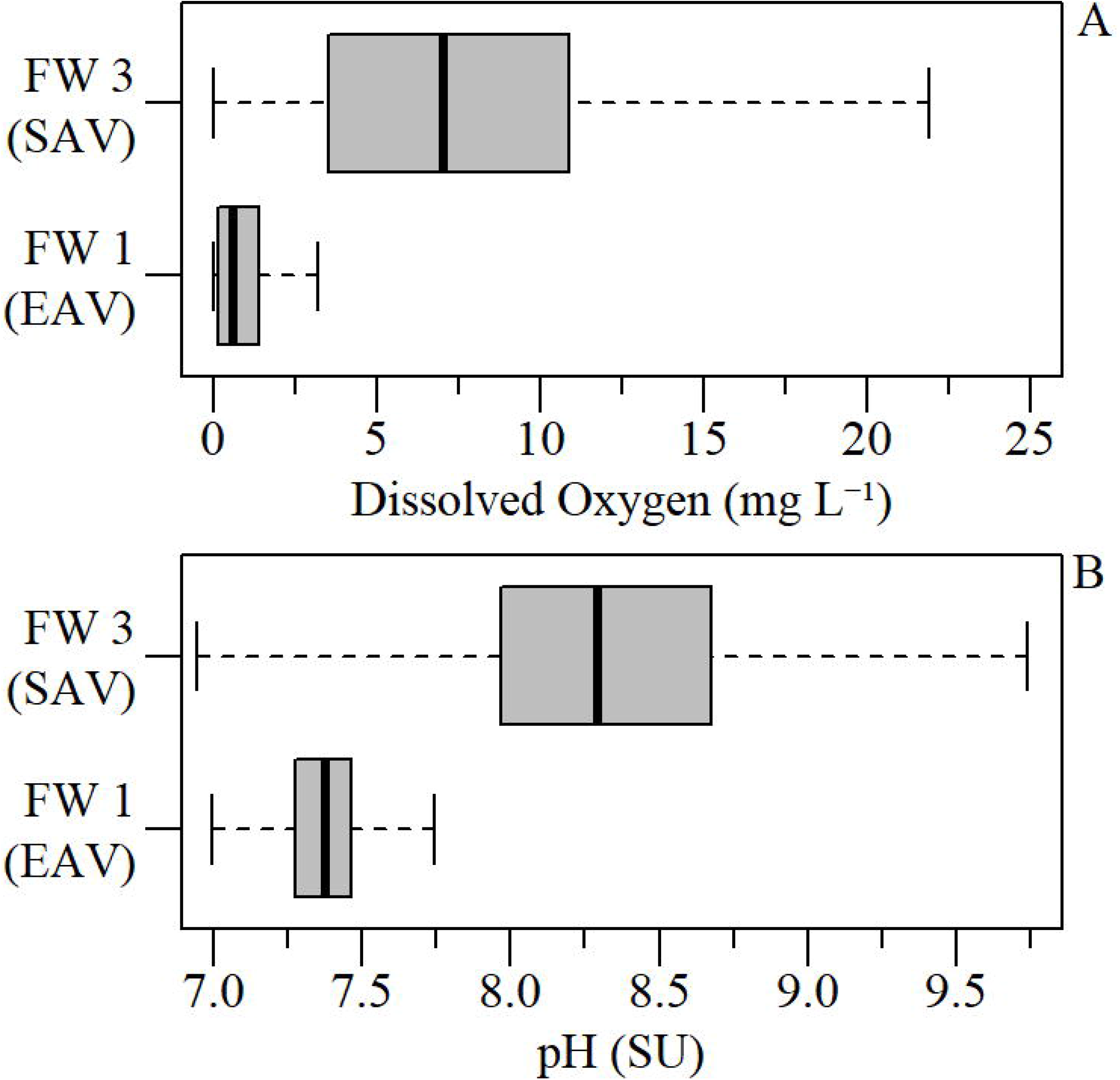
Dissolved oxygen (A) and pH (B) across all sites along the flow-way (FW) 1 and FW 3 transects within Stormwater Treatment Area 2 measured during this study. Dominate vegetation of FW 1 and 3 is emergent aquatic vegetation (EAV) and submerged aquatic vegetation (SAV), respectively.

**Table 1.** Summary statistics for parameters and matrices used in this study from samples collected along flow-way (FW) 1 and 3 within Stormwater Treatment Area-2. Summary statistics expressed as mean ± standard error (sample size). Matrices include surface water, flocculent material and recently accreted soil.

| Matrix | Parameter | Units | FW1 | FW3 |
| --- | --- | --- | --- | --- |
| Surface Water | Dissolved Oxygen (DO)<sup>1</sup> | mg L<sup>-1</sup> | 1.02 ± 0.01 (13447) | 7.89 ± 0.04 (19543) |
| | pH<sup>1</sup> | SU | 7.35 ± 0.001 (17685) | 8.38 ± 0.004 (20281) |
| | Temperature<sup>1</sup> | °C | 27.2 ± 0.02 (17129) | 26.3 ± 0.03 (20288) |
| | Specific Conductivity<sup>1</sup> | μS cm<sup>-1</sup> | 842.4 ± 1.3 (17280) | 941.9 ± 1.4 (20290) |
| | Turbidity<sup>1</sup> | NTU | 4.0 ± 0.2 (16779) | 4.5 ± 0.06 (17004) |
| | Gross Primary Productivity (GPP)<sup>2</sup> | g O<sub>2</sub> m<sup>-2</sup> d<sup>-1</sup> | 0.83 ± 0.03 (750) | 6.46 ± 0.16 (805) |
| | Ecosystem Respiration (ER)<sup>2</sup> | g O<sub>2</sub> m<sup>-2</sup> d<sup>-1</sup> | -0.84 ± 0.03 (750) | -6.72 ± 0.17 (805) |
| | Net Aquatic Productivity (NAP)<sup>2</sup> | g O<sub>2</sub> m<sup>-2</sup> d<sup>-1</sup> | -0.02 ± 0.007 (750) | -0.27 ± 0.04 (805) |
| | Total Phosphorus (TP)<sup>3</sup> | μg L<sup>-1</sup> | 58.3 ± 5.3 (154) | 35.1 ± 1.9 (229) |
| | Dissolved Organic Carbon (DOC)<sup>3</sup> | mg L<sup>-1</sup> | 27.2 ± 0.5 (143) | 30.9 ± 0.2 (237) |
| | Chlorophyll *a*<sup>3</sup> | μg L<sup>-1</sup> | 5.0 ± 0.5 (105) | 12.2 ± 1.3 (132) |
| Floc | Total Carbon (TC) | g kg<sup>-1</sup> | 406 ± 9 (34) | 200 ± 5 (35) |
| | Bulk Density (BD) | g cm<sup>-3</sup> | 0.03 ± 0.003 (37) | 0.12 ± 0.007 (37) |
| | Loss-on-ignition (LOI) | % | 76.4 ± 2.1 (34) | 22.7 ± 1.2 (35) |
| | Carbon Storage<sup>4</sup> | g m<sup>-2</sup> | 603 ± 37 (34) | 1918 ± 162 (30) |
| Soil | Total Carbon (TC) | g kg<sup>-1</sup> | 445 ± 7 (51) | 351 ± 16 (51) |
| | Bulk Density (BD) | g cm<sup>-3</sup> | 0.08 ± 0.004 (37) | 0.22 ± 0.01 (37) |
| | Loss-on-ignition (LOI) | % | 81.2 ± 1.4 (51) | 54.2 ± 3.3 (51) |
| | Carbon Storage<sup>4</sup> | g m<sup>-2</sup> | 766 ± 84 (34) | 1867 ± 202 (33) |

<sup>1</sup> Data from 15-min/30-min logged data using water quality sondes deployed during flow events.
<sup>2</sup> Estimated from 15-min/30-min logged data using water quality sondes and nearby weather monitoring locations during flow events.
<sup>3</sup> Data collected via weekly grab samples and QA/QC screened accordingly.
<sup>4</sup> Carbon storage estimated for samples using equation 4.

In aquatic ecosystems, it is generally assumed that the net balance of DO is governed by metabolism. However, as discussed in the context of lake ecosystems changes in DO can also be influenced by interaction with DO depleted groundwater, physicochemical interactions, photorespiration of dissolved organic matter (Osborne et al., *In Prep.*) and exchange with the atmosphere (Cole et al. 2000; Hanson et al. 2003). Generally DO concentrations are depressed in wetland ecosystems, including oligotrophic wetlands due to the mineralization of allochthonous organic matter resulting in net heterotrophic conditions (Hagerthey et al. 2010).

In net heterotrophic lakes it was determined that groundwater driven DO flux was minor and did not significantly effect estimates of NAP (Cole et al. 2000). Generally, groundwater is low in DO but can contribute to the total C budget of an aquatic ecosystem by providing a significant input of DIC as demonstrated by Staehr et al. (2010). Additionally, Staehr et al. (2010) indicated that allochthonous inputs of dissolved organic matter (DOM) via streams are generally the dominate source of C to lake ecosystems, is slowly degraded relative to autochthonous DOM and in dystrophic lakes accounts for the majority of C mineralization in the system. In the STAs, seepage can vary between FWs because of the underlying geology, conductivity of levee material and difference in water levels between compartments. Specific to the FWs in this study, average annual seepage account for ~1% of the total volume entering the FW while seepage out accounts for ~5% of the outflow water budget (Zhao and Piccone 2018). Given this minor contribution to the system and the relatively high C storages in the soil (Table 1), C from groundwater/seepage is a minor contribution to the overall C pool available for mineralization within these FWs.

### Aquatic Metabolism

Rates of GPP, NEP and R varied between the FWs, generally with FW 3 exhibited an order of magnitude difference (Fig 4 and S3). Across the entire study and between the two FWs, GPP ranged from −1.2 to 22.5 g O_2_ m^−2^ d^−1^, ER ranged from −32.3 to 2.0 g O_2_ m^−2^ d^−1^ and NAP ranged from −9.8 to 3.3 g O_2_ m^−2^ d^−1^. The range of these values reported in this study are consistent with a prior study within the Everglades ecosystem (Hagerthey et al. 2010). Hagerthey et al. (2010) evaluated aquatic metabolism of several ecosystems, with a wide range of stressors, some analogous to the ecosystems assessed in this study. The two ecosystem types that closely resembles the community assemblage of the systems in this study was an open-water slough dominated by SAV with mineral soils similar to FW 3 and a dense monoculture of *Typha* with organic soils comparable to FW 1. For these ecosystems, Hagerthey et al. (2010) reported higher DO concentrations/% saturation, greater GPP and NAP and lower water column TP concentrations for the SAV dominated open-water slough (FW 3 analogous) relative to the EAV dominated ecosystem (FW 1 analogue).

**Figure 4.**
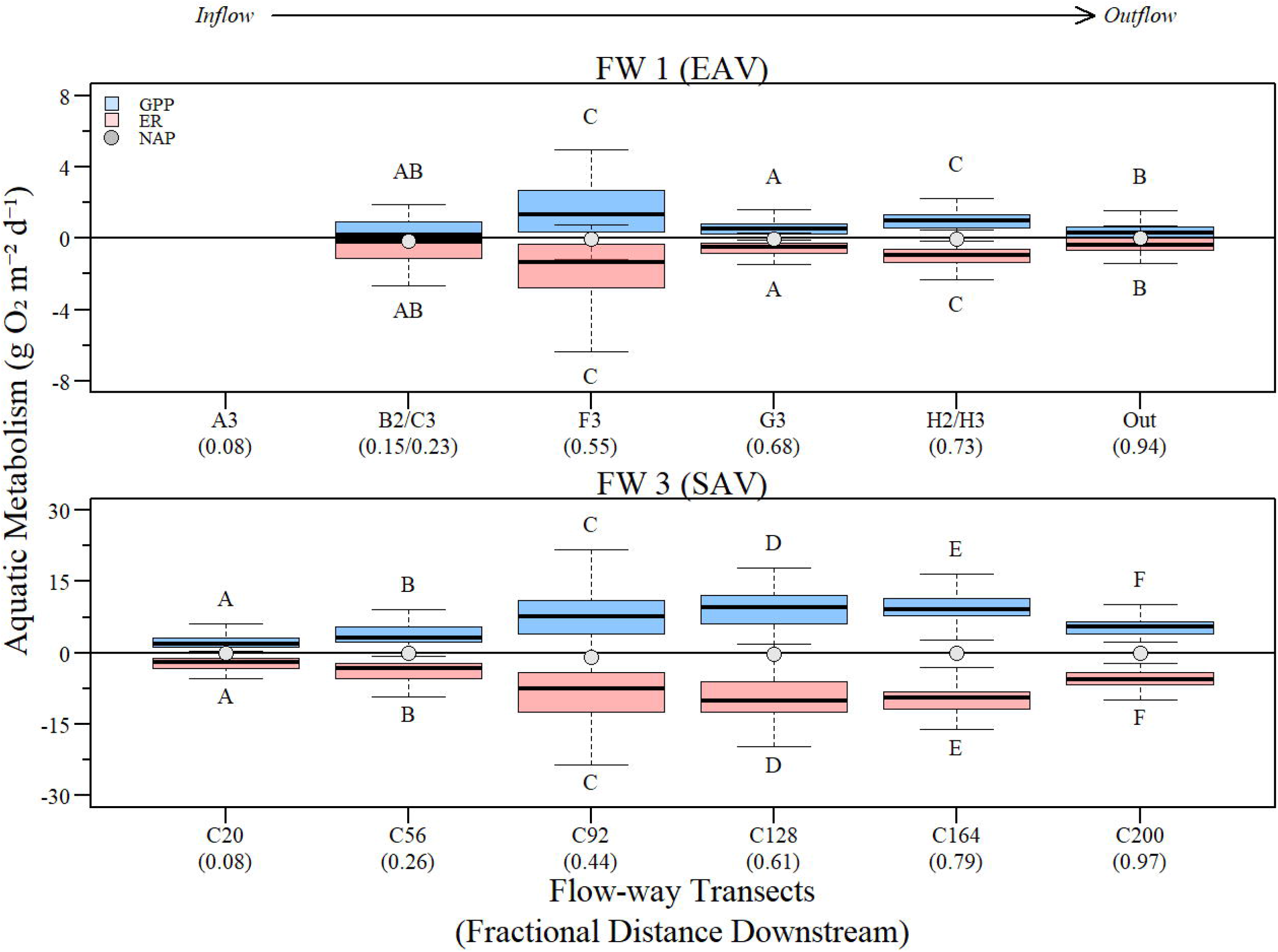
Boxplot of daily gross primary productivity (GPP), ecosystem respiration (ER) and mean net aquatic productivity (NAP) along the Stormwater Treatment Area 2 flow-way 1 transect, moving from inflow to outflow with site and fractional distance downstream (0.0 = inflow; 1.0 = outflow). Letters indicate significant differences between sites for GPP and ER according to Dunn’s Multiple Comparisons.

In this study we report slightly negative GPP and slightly positive ER values, as discussed by Caffrey et al. (2014) some studies consider these values anomalous and are typically discarded. Factors that contribute to these potential anomalous GPP and ER values could be underestimation of re-aeration coefficients, modeling artifacts or period of low metabolism below the limit of detection (Caffrey et al. 2014; Demars et al. 2015; Hornbach et al. 2017). Re-aeration coefficients can either be estimated *in-situ* via tracer injection methods or empirically estimated using meteorological and water temperature (this study; Thébault et al. 2008; Caffrey et al. 2014). Estimating re-aeration coefficients with tracer based studies relies on a large quantity of data to successfully temporally scale across a lager study (Demars et al. 2015). Volumetric re-aeration as implemented by Thébault et al. (2008) and this study is more robust to fine scale temporal estimation given it uses meteorological variables (wind speed, barometric pressure, air temperature and water temperature) and water column depth to estimates the variable. Given the robustness of the estimate, GPP and ER calculations are relatively insensitive to uncertainty in estimation of volumetric re-aeration (Fig S5). Moreover, under or over estimation of re-aeration coefficients would only amplify changes in GPP or ER not change the sign (Demars et al. 2015 and Fig S5). It is more likely that these presumably anomalous values indicate periods of low metabolism below the effective level of detection, given the extreme differences in DO concentrations over the diel cycle and prolonged periods (several hours) of anoxic conditions.

Measurement of aquatic metabolism is a fundamental metric of an ecosystems function and represents an integrative process of OM production (via GPP) and consumption (via ER) via aerobic respiration (Odum 1956; Staehr et al. 2012; Hoellein et al. 2013). These metrics of OM processing also reflects system-level responses to external perturbations including allochthonous input of OM and growth limiting nutrients. In the long term (i.e. annual, decadal, etc.) production and consumption of OM is expected to balance, however metabolic responses over shorter periods (i.e. daily, seasonal, etc.) identify characteristic effects of disturbance (Odum 1956). These short-term responses can manifest as relative changes in GPP and ER magnitude. Disturbances such an episodic surge in stream flow and wind-driven events have the potential to cause sediment resuspension resulting in ecosystem level changes in GPP and ER. Additionally, high flow events deliver pulses of inorganic nutrients, DOC and suspended sediment depending on the system. These inputs can induce both positive and negative effects on GPP and ER (Flöder and Sommer 1999; Hanson et al. 2008; Staehr et al. 2010, 2012; Tsai et al. 2011). In general, the Everglades STAs are hydrologically dynamic (i.e. pulsed) due to the intended purpose of the system to treat stormwater run-off from the upstream watershed (Juston and DeBusk 2006; Juston and Kadlec 2019). Along with these pulses the system receives allochthonous nutrients and C which has the potential to influence GPP and ER. Meanwhile, vegetation ecophysiology and response to these changes in hydrodynamic and nutrient condition combined with *in-situ* changes in water level, water transparency (i.e. attenuation), etc. can also have profound effects on system wide GPP and ER.

In longitudinally linked systems such as streams and flowing wetlands downstream ecological processes are linked to upstream areas though longitudinal transport concurrent with changes in energy and material inputs. This longitudinal linkage is the foundation of stream nutrient spiraling concept and biogeochemical processing of nutrients in lotic ecosystems (Newbold 1987; Webster 2007). In river ecosystems, transport displaces autochthonous and allochthonous OM downstream producing a metabolism gradient along the river continuum driven by OM recalcitrance, transportability and the mineralization of OM-bound nutrients (Webster 2007). In the STAs, nutrients, material and energy are transported, albeit much slower than stream and river systems longitudinally along the wetland where autochthonous and allochthonous OM and nutrients are metabolized ultimately storing C and material along the FW. In this highly managed system allochthones inputs are generally isolated to inflow regions and transported downstream. Along the two FWs in this study GPP (FW 1: χ^2^=102.1, df=4,ρ<0.01; FW 3: χ^2^=368.2, df=5, ρ<0.01) and ER (FW 1: χ^2^=98.5, df=4,ρ<0.01; FW 3: χ^2^=373.0, df=5, ρ<0.01) were significantly different between sites with an order of magnitude difference between FWs (Fig 4). In FW 3, all sites/FW position significantly differed with GPP and ER reaching a maximum near the FW mid-point with a slight decline in the back two-thirds of the FW (Fig 4). This trend in GPP and ER observed in FW 3 is opposite to the P concentration gradient observed during this study (UF-WBL 2017) and elsewhere (Juston and Kadlec 2019). Trends in GPP, ER and P reductions mirror changes in pH (Fig 3) along the FW suggesting in addition to biotic retention of P, abiotic processes are involved with P reductions along the FW.

### Loading Conditions

During this study, a total of six prescribed/managed flow events occurred between August 10^th^, 2015 and July 31^st^, 2017 with events ranging from 35 to 63 days in FWs 1 and 3 within STA-2. During the flow events, daily HLR ranged between 0 (no inflow) to a peak loading of 18.1 cm d^−1^ with FW 3 receiving a relatively higher mean HLR of 3.4 ± 0.3 cm d^−1^ (Mean ± SE), compared to 2.2 ± 0.4 cm d^−1^. These FW specific daily HLR rates during experimental flow events correspond to STA wide long term HLR values reported by Chen et al. (2015) of 3.3 ± 0.8 cm d^−1^. Observed daily PLR values ranged from 0 (no loading) to 38.3 mg m^−2^ d^−1^ with FW 1 receiving a higher relative loading rate 3.4 ± 0.7 mg m^−2^ d^−1^ compared to FW 3 with a mean PLR of 2.1 ± 0.2 mg m^−2^ d^−1^ (Supplemental Material; Table S2 and Fig S1). These PLR rates are well within the rates observed by Chen et al. (2015) of approximately 2.6 mg m^−2^ d^−1^ (converted from 0.96 g m^−2^ year^−1^). The daily HLR and PLR observed during this study was consistent with historic daily operational loading rates experienced for these FWs in the recent past (Fig S1) and to date (Chen et al. 2015; Chimney 2019). Mean HLR values observed during the study occurred at 0.56 and 0.71, respectively for FW 1 and FW 3 along the FW HLR cumulative duration curves (Fig S1 and S2). Meanwhile, mean PLR values observed during the study relative to the recent historical time-series occurred at 0.71 for both FW 1 and FW 3 along their respective PLR CDF curves despite FW 3 having a higher maximum PLR and steeper CDF curve (Fig S1 and S2). These results suggest a similar P loading regime relative to the period of record despite slightly different HLR. The difference in HLRs between FWs is presumably driven by operational targets or benchmarks such as target water depths, loading rates, water management and best professional judgement of water managers. Moreover, these FWs and larger STA is part of an integrated water management system to provide flows to the downstream Everglades and as such is at the whim of climatic variability and event based operations to some extent (Pietro 2012).

Aquatic GPP in some ecosystems such as rivers and stream can be stimulated by periodic hydrologic pulses by altering the benthic habitat potentially suppressing predators, removing organic waste material or clearing material to create fresh substrate for colonization of benthic algae (Mosisch and Bunn 1997). Alternatively, high velocity flows can have detrimental effects to benthic communities by dislodging material and exporting biomass form the system and excessive inundation of producers (Stevenson et al. 1996; Mosisch and Bunn 1997). Moreover, O’Donnell and Hotchkiss (2019) demonstrated in that in stream ecosystems that changing flow conditions interact with biotic ad chemical processes resulting in divergent metabolic responses. In this study, HLR was significantly correlated with spatially averaged GPP (r_s_=0.40, ρ<0.01), ER (r_s_= −0.42, ρ<0.01) and NAP (r_s_= −0.17, ρ<0.05) across FW 3. Meanwhile, spatially averaged GPP (r_s_=0.06, ρ=0.46), ER (r_s_= −0.07, ρ=0.41) and NAP (r_s_= −0.17, ρ<0.05) were not significantly correlated across FW 1 suggesting that ecosystem metabolism is controlled differently across these systems and that loading/transport alone is not the only driver of ecosystem metabolism much like some river ecosystems (Webster 2007).

### Soil carbon linkage

Aquatic metabolism is the short-term measure of GPP and C sequestration within aquatic ecosystems. Variability in irradiance and temperature can cause dynamic changes in the rates of photosynthesis and community respiration within a given ecosystem can cause short term, and at times rapid changes in rates of GPP, ER and NAP (Staehr and Sand-Jensen 2007). As Hagerthey et al. (2010) suggests the patterns of NAP in the downstream Everglades is driven largely by a large standing crop (i.e. biomass) combined with high productivity of vegetation contributing autochthonous C inputs. These inputs from macrophytes are the result of complex interactions of abiotic and biotic processes in the ambient environment through pulsed nutrients and hydrological inputs over time (Ewe et al. 2006). The fate of this autochthonous C inputs in the form of leaflitter or senescent material accumulates as floc and eventually becomes incorporated into the soil. Along this decay continuum, vertebrates (detritovores)(Blanco et al. 2004), invertebrate (shredders)(Bohman and Tranvik 2001) and microbial communities (Yarwood 2018) contribute to the breakdown of this C contributing to the DOM pool (i.e. water column DOC) further stimulating aquatic metabolism leaving the remaining to be buried in recently accreted soil (Fig 5). Like the downstream Everglades, the STAs are use the natural communities of the Everglades (i.e. cattail, sawgrass, chara, etc.) to facilitate the removal of nutrients. The STAs have a large standing crop and highly productive vegetation that are subsidized by nutrients frequently as water is treated in the system producing large quantities of autochthonous C.

**Fig 5.**
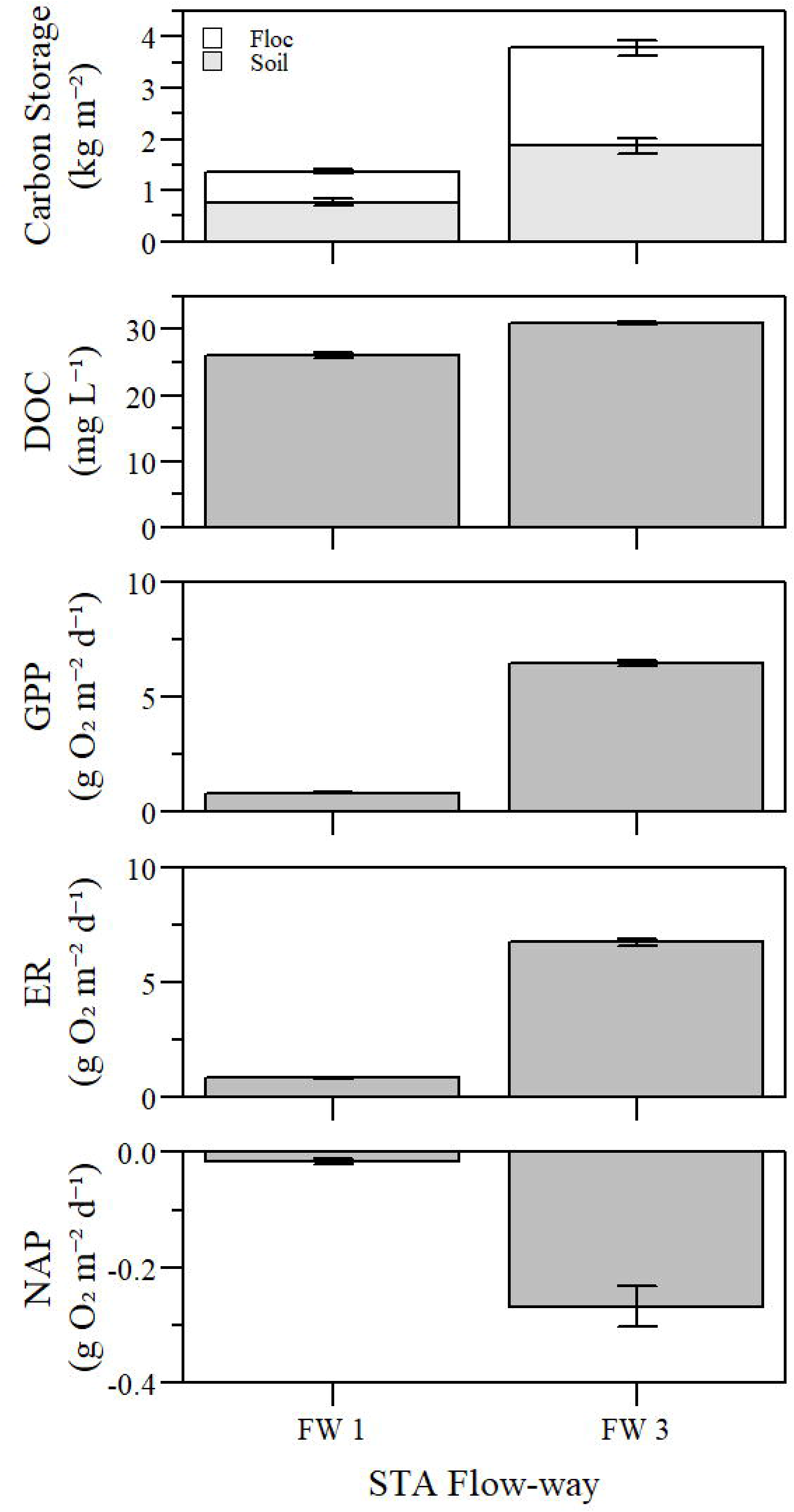
Flow-way (FW) mean (±SE) soil carbon storage, water column dissolved organic carbon (DOC), gross primary production (GPP), ecosystem respiration (ER) and net aquatic productivity (NAP) observed during this study along each FW transect.

Due to the limited deployments of the sondes and the timing of the soil sampling no statistical analysis was conducted to evaluate soil C storage and aquatic metabolism, however the qualitative relationship between soil C storage and that of aquatic productivity is striking. On average, soil C storage and surface DOC concentrations are greater in the FW 3 where GPP and ER are greatest (Fig 5). Given the extreme pH variability in FW 3 (Fig 3) and the ionic characteristics of the water column (UF-WBL 2017), the proportion of dissolved organic and inorganic C could differ significantly between FWs. While FW 3 C storages are qualitatively greater than FW 1, the composition of that total C could differ between FWs. Based on high soil calcium concentrations (Julian et al. In Press) and low LOI values (Table 1), generally FW 3 soils have high inorganic C storage (UF-WBL 2017) driven by the removal calcium carbonate via SAV photosynthesis and respiration (Dierberg et al. 2002) while the C stored in FW 1 is predominately organic (Table 1). Both FWs store significant amounts of C with FW 3 storing a larger quantity, but the composition and quality of the C could differ and therefore influencing other biogeochemical cycles.

Despite difference in soil C composition, both FWs have large storages of C in floc and soil. The trend in C storages within FW mirrors that of aquatic metabolism estimates. While storages do not directly equate to productivity it is an indicator of ecosystem production and biogeochemical cycling of nutrients and matter. Notwithstanding the extreme variability in aquatic productivity across and between FWs show that both are net accreting systems (Bhomia et al. 2015) and are generally heterotrophic with respect to water column metabolism (Fig 6) with both FWs GPP to ER ratios being significantly less than one (FW 1: V=106430, ρ<0.01; FW 3: V=123510, ρ<0.01). Much like in most lake ecosystems, the STAs in this study experience a GPP to ER ratio of less than one indicates that the is ER is subsidized by allochthonous OM (Cole et al. 2000). This OM subsidy could explain the change in ER in GPP in FW 3 at high ER rates where the data falls below the 1:1 line (Fig 6). Meanwhile, GPP is frequently limited by the supply of plant-limiting nutrients in lake ecosystems (Elser et al. 1990) and is the case with shallow wetlands with respect to aquatic and periphyton biomass-specific productivity (McCormick et al. 1998; Hagerthey et al. 2010). In the case of the STAs should we expect that GPP and P concentrations be indicative of strong P-removal processes?

**Figure 6.**
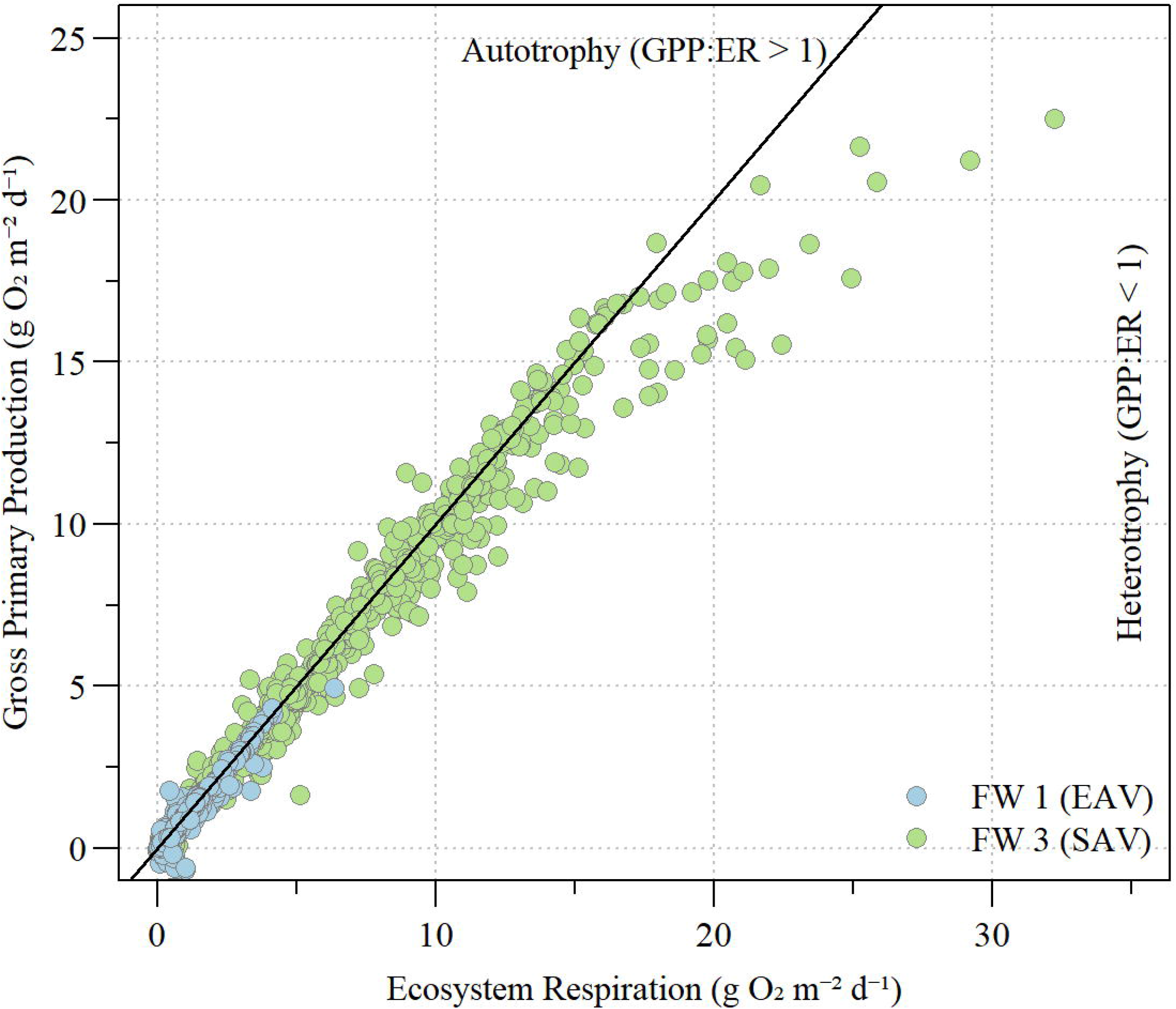
Daily water column measurements of metabolism in Stormwater Treatment Area 2, flow-way (FW) 1 and 3 across the study period. The solid line indicates 1:1 vale of Gross Primary Production (GPP) to Ecosystem Respiration (ER).

### Aquatic productivity and Water Quality

The Everglades STAs are unique treatment wetland ecosystems constructed to reduce water column P concentrations to protect the downstream oligotrophic Everglades (Chen et al. 2015). Leveraging ecosystem and biological functionality of wetland ecosystems this reduction in P is typically achieved. As stated above, the supply of limiting nutrients can have a significant influence on GPP and thus NAP. Moreover, habitat dynamics, macrophyte canopy shading and legacy enrichment can all play a significant roles in aquatic ecosystem metabolism (Hagerthey et al. 2010). In the case of our study, each ecosystem (i.e. EAV and SAV) were evaluated in relative isolation from each other with notable differences in NAP (Fig 4) and reflected in water column and soil C dynamics (Fig 5). Moreover, concurrent with greater GPP and ER values, FW 3 also experienced a greater diel pH relationship. In SAV systems, a P-removal pathway is driven by photosynthetically driven mediation of water column pH resulting in concomitant precipitation of P out of solution as Ca-P compounds stimulating particulate P production (Dierberg et al. 2002; White et al. 2006; Juston and Kadlec 2019).

In lake ecosystems, GPP is typically positively correlated with limiting nutrient concentrations (Hanson et al. 2003). However in the downstream Everglades marsh, Hagerthey et al. (2010) observed a negative correlation between TP and GPP suggesting the casual mechanism for this relationship is regulated by light limitations caused by enhanced standing stock of emergent vegetation, periphyton and algal biomass. In this study, GPP was inversely related to water column TP concentrations in FW 3 (r_s_ = −0.20; ρ<0.01) but not in FW 1 (r_s_ = 0.02; ρ=0.81). The lack of a correlation observed in FW 1 could be the result of macrophyte shading reducing light penetration to and through the water column depressing aquatic GPP as suggested by Hagerthey et al. (2010), however GPP was inversely correlated with water column chlorophyll *a* concentration (r_s_ = −0.25; ρ<0.05) in FW 1 and borderline significant in FW 3 (r_s_ = −0.18; ρ=0.05). While both FWs are negatively correlated with daily mean turbidity (FW 1: r_s_ = −0.26; ρ<0.01; FW 3: r_s_ = −0.53; ρ<0.01). The lack correlation between GPP and TP within FW 1 could also be due to data limitation relative to the experimental flow event. Despite this limitation it could be suggested that GPP is regulated by SAV productivity in the case of FW 3 and possibly algal productivity in FW 1 (Stanley et al. 2003). In addition to direct biotic regulation of GPP, other factors such as turbidity and limiting nutrients which drive light limitations and periphyton productivity can also influence aquatic metabolism dynamics within these wetlands systems consistent with prior studies (McCormick et al. 1998; Hagerthey et al. 2010). However, what is not directly obvious is if along the FW a detectable change point between the water column TP and GPP is obvious and directly related to STA function (Fig 7). In addition to water column dynamics, these FWs also have an underlying soil nutrient gradient (Osborne et al. *In Prep*) which could also influence short-term aquatic metabolism dynamics. More study is needed to evaluate the response of ecosystem metabolism relative to P-cycling, especially in the lower reaches these FWs where TP concentrations are in the low-P domain and GPP in the case of FW 3 begins to decline.

**Figure 7.**
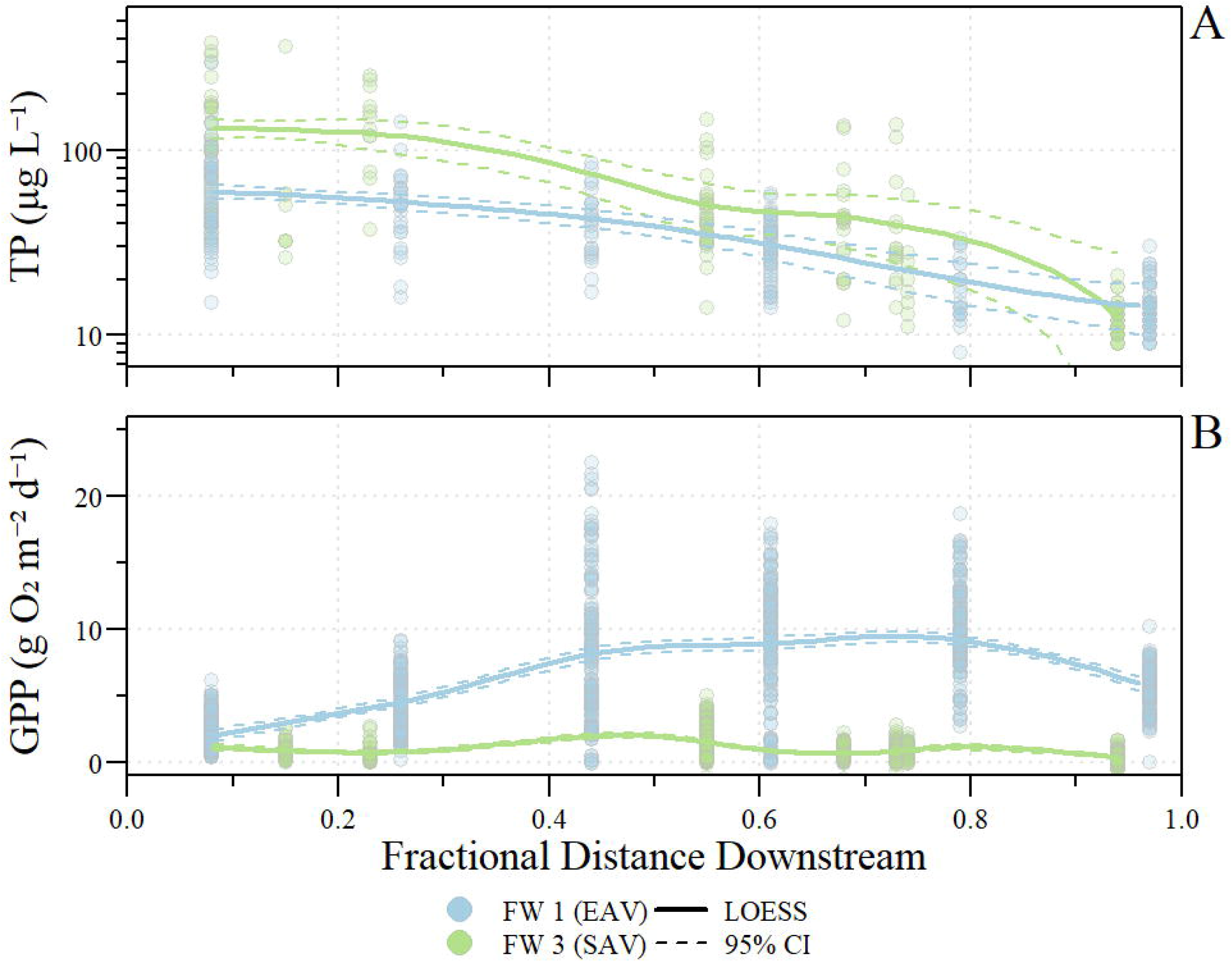
A) Weekly total phosphorus (TP) water column grab samples and B) daily estimated gross primary productivity (GPP) during experimental flow events by fractional distance downstream along stormwater treatment area 2, flow-way (FW) 1 and 3. A fractional distance of zero (0) is the inflow to the FW and fraction distance of one (1) is outflow of the FW. Locally estimated scatterplot smoothing (LOESS) regression ± 95% confidence interval was estimated using all data for each FW along the transect.

### Conclusions and future considerations

Patterns of O_2_ dynamics and thus aquatic metabolism observed in this study vary spatially (along FWs) and temporally (diel to days) controlled by different effects related biological, physical and chemical processes. This study supports conclusions by Hagerthey et al. (2010) that aboveground biota regulate aquatic metabolism and O_2_ dynamics in shallow wetland ecosystems by focusing on two STA FWs with two distinct dominate vegetative communities. This study suggests that ecosystem metabolism is controlled differently across FWs with varying levels of response to loading/transport and water column attributes resulting in differences OM and C turnover as indicated by differences in soil C and water column DOC concentrations between FWs. As observed by Hagerthey et al. (2010) and this study dense emergent macrophytes (EAV) limits O_2_ exchange with the atmosphere and limits O_2_ availability combined with a high sediment oxygen demand results in predominately anaerobic respiration reducing C turnover and lowering overall aquatic productivity. Conversely SAV facilitates aerobic conditions by aerating the water column resulting in large diel swings in O_2_ resulting in greater GPP and C turnover. In addition to differences due dominate vegetative communities, aquatic metabolism in each FW responded differently relative to HLR, water column P, and chlorophyll *a* concentration. This study was an initial investigation into aquatic metabolism in a treatment wetland system designed to remove P from stormwater runoff prior to entering the Everglades ecosystem. This system is also a strategy to protect and preserve the downstream Everglades by improve nutrient conditions entering the ecosystem. Understanding the limitation of these system to removing P is essential in achieving and maintaining necessary levels of treatment. Additional studies in aquatic metabolism and overall system productivity is needed to provide further understanding in how productivity and respiration relate to treatment performance.

## Acknowledgements

We would like to thank SFWMD and UF Wetland Biogeochemistry Laboratory staff members for providing the data used in this analysis. We would also like to thank Jill King, Odi Villapando, Cassondra Armstrong, Tom Jones and the anonymous peer reviewer(s) and editor(s) for their efforts and constructive review of this manuscript.

## Authors’ Contributions

PJ performed data analyses including necessary calculations and statistical analyses and wrote the manuscript. TZO was involved with soil sample collection and writing of the manuscript.

## References

Beck MW (2016) SWMPr: Retrieving, Organizing, and Analyszing Estuary Monitoring Data,. CRAN R-Project

Bhomia RK, Inglett PW, Reddy KR (2015) Soil and phosphorus accretion rates in sub-tropical wetlands: Everglades Stormwater Treatment Areas as a case example. Science of The Total Environment 533:297–306. https://doi.org/10.1016/j.scitotenv.2015.06.115

Billett MF, Moore TR (2008) Supersaturation and evasion of CO2 and CH4 in surface waters at Mer Bleue peatland, Canada. Hydrol Process 22:2044–2054. https://doi.org/10.1002/hyp.6805

Blanco S, Romo S, Villena M-J (2004) Experimental Study on the Diet of Mosquitofish (Gambusia holbrooki) under Different Ecological Conditions in a Shallow Lake. International Review of Hydrobiology 89:250–262. https://doi.org/10.1002/iroh.200310684

Bohman IM, Tranvik LJ (2001) The effects of shredding invertebrates on the transfer of organic carbon from littoral leaf litter to water-column bacteria. Aquatic Ecology 35:43–50. https://doi.org/10.1023/A:1011425905036

Bruland GL, Osborne TZ, Reddy KR, et al (2007) Recent Changes in Soil Total Phosphorus in the Everglades: Water Conservation Area 3. Environmental Monitoring and Assessment 129:379–395. https://doi.org/10.1007/s10661-006-9371-x

Caffrey JM, Murrell MC, Amacker KS, et al (2014) Seasonal and Inter-annual Patterns in Primary Production, Respiration, and Net Ecosystem Metabolism in Three Estuaries in the Northeast Gulf of Mexico. Estuaries and Coasts 37:222–241. https://doi.org/10.1007/s12237-013-9701-5

Caraco N, Cole J, Findlay S, Wigand C (2006) Vascular Plants as Engineers of Oxygen in Aquatic Systems. BioScience 56:219–225. https://doi.org/10.1641/0006-3568(2006)056[0219:VPAEOO]2.0.CO;2

Caraco NF, Cole JJ (2002) Contrasting Impacts of a Native and Alien Macrophyte on Dissolved Oxygen in a Large River. Ecological Applications 12:1496–1509. https://doi.org/10.1890/1051-0761(2002)012[1496:CIOANA]2.0.CO;2

Carignan R, Planas D, Vis C (2000) Planktonic production and respiration in oligotrophic Shield lakes. Limnology and Oceanography 45:189–199. https://doi.org/10.4319/lo.2000.45.1.0189

Carpenter SR, Lodge DM (1986) Effects of submersed macrophytes on ecosystem processes. Aquatic Botany 26:341–370. https://doi.org/10.1016/0304-3770(86)90031-8

Carpenter SR, Pace ML (1997) Dystrophy and Eutrophy in Lake Ecosystems: Implications of Fluctuating Inputs. Oikos 78:3–14. https://doi.org/10.2307/3545794

Chen H, Ivanoff D, Pietro K (2015) Long-term phosphorus removal in the Everglades stormwater treatment areas of South Florida in the United States. Ecological Engineering 79:158–168. https://doi.org/10.1016/j.ecoleng.2014.12.012

Chimney M (2019) Performance of the Everglades Stormwater Treatment Areas. In: 2019 South Florida Environmental Report. South Florida Water Management District, West Palm Beach, FL

Chimney MJ, Goforth G (2001) Environmental impacts to the Everglades ecosystem: a historical perspective and restoration strategies. Water Science & Technology 44:93–100

Cole JJ, Pace ML, Carpenter SR, Kitchell JF (2000) Persistence of net heterotrophy in lakes during nutrient addition and food web manipulations. Limnol Oceanogr 45:1718–1730. https://doi.org/10.4319/lo.2000.45.8.1718

Corstanje R, Grafius DR, Zawadzka J, et al (2016) A datamining approach to identifying spatial patterns of phosphorus forms in the Stormwater Treatment Areas in the Everglades, US. Ecological Engineering 97:567–576. https://doi.org/10.1016/j.ecoleng.2016.10.003

DeBusk TA, Grace KA, Dierberg FE, et al (2004) An investigation of the limits of phosphorus removal in wetlands: a mesocosm study of a shallow periphyton-dominated treatment system. Ecological Engineering 23:1–14. https://doi.org/10.1016/j.ecoleng.2004.06.009

del Giorgio PA, Peters RH (1994) Patterns in planktonic P: R ratios in lakes: Influence of lake trophy and dissolved organic carbon. Limnol Oceanogr 39:772–787

Demars BOL, Thompson J, Manson JR (2015) Stream metabolism and the open diel oxygen method: Principles, practice, and perspectives: Problems in stream metabolism studies. Limnol Oceanogr Methods 13:356–374. https://doi.org/10.1002/lom3.10030

Dierberg FE, DeBusk TA, Jackson SD, et al (2002) Submerged aquatic vegetation-based treatment wetlands for removing phosphorus from agricultural runoff: response to hydraulic and nutrient loading. Water research 36:1409–1422

Dinno A (2015) Dunn’s test of multiple comparisons using rank sums. CRAN R-Project

Elser JJ, Marzolf ER, Goldman CR (1990) Phosphorus and Nitrogen Limitation of Phytoplankton Growth in the Freshwaters of North America: A Review and Critique of Experimental Enrichments. Can J Fish Aquat Sci 47:1468–1477. https://doi.org/10.1139/f90-165

Ewe SML, Gaiser EE, Childers DL, et al (2006) Spatial and temporal patterns of aboveground net primary productivity (ANPP) along two freshwater-estuarine transects in the Florida Coastal Everglades. Hydrobiologia 569:459–474. https://doi.org/10.1007/s10750-006-0149-5

Flöder S, Sommer U (1999) Diversity in planktonic communities: An experimental test of the intermediate disturbance hypothesis. Limnol Oceanogr 44:1114–1119. https://doi.org/10.4319/lo.1999.44.4.1114

Freeman C, Fenner N, Ostle NJ, et al (2004) Export of dissolved organic carbon from peatlands under elevated carbon dioxide levels. Nature 430:195–198

Hagerthey SE, Cole JJ, Kilbane D (2010) Aquatic metabolism in the Everglades: Dominance of water column heterotrophy. Limnology and Oceanography 55:653–666

Hanson PC, Bade DL, Carpenter SR, Kratz TK (2003) Lake metabolism: Relationships with dissolved organic carbon and phosphorus. Limnol Oceanogr 48:1112–1119. https://doi.org/10.4319/lo.2003.48.3.1112

Hanson PC, Carpenter SR, Kimura N, et al (2008) Evaluation of metabolism models for free-water dissolved oxygen methods in lakes. Limnology and Oceanography: Methods 6:454–465. https://doi.org/10.4319/lom.2008.6.454

Harris W (2011) Mineral Distribution and Weathering in the Greater Everglades: Implications for Restoration. Critical Reviews in Environmental Science and Technology 41:4–27. https://doi.org/10.1080/10643389.2010.531191

Hoellein TJ, Bruesewitz DA, Richardson DC (2013) Revisiting Odum (1956): A synthesis of aquatic ecosystem metabolism. Limnol Oceanogr 58:2089–2100. https://doi.org/10.4319/lo.2013.58.6.2089

Hornbach DJ, Hove MC, Ensley-Field MW, et al (2017) Comparison of ecosystem processes in a woodland and prairie pond with different hydroperiods. Journal of Freshwater Ecology 32:675–695. https://doi.org/10.1080/02705060.2017.1393468

Hotchkiss ER, Sadro S, Hanson PC (2018) Toward a more integrative perspective on carbon metabolism across lentic and lotic inland waters. Limnology and Oceanography Letters 3:57–63. https://doi.org/10.1002/lol2.10081

Julian P, Gerber S, Bhomia RK, et al (In Press) Evaluation of nutrient stoichiometric relationships among ecosystem compartments of a subtropical treatment wetland. Do we have “Redfield wetlands”? Ecological Processes

Julian P, Gerber S, Bhomia RK, et al (2019) Evaluation of nutrient stoichiometric relationships among ecosystem compartments of a subtropical treatment wetland. Do we have “Redfield wetlands”? Ecol Process 8:20. https://doi.org/10.1186/s13717-019-0172-x

Julian P, Gerber S, Wright AL, et al (2017) Carbon pool trends and dynamics within a subtropical peatland during long-term restoration. Ecol Process 6:43–57. https://doi.org/10.1186/s13717-017-0110-8

Juston J, DeBusk TA (2006) Phosphorus mass load and outflow concentration relationships in stormwater treatment areas for Everglades restoration. Ecological Engineering 26:206–223. https://doi.org/10.1016/j.ecoleng.2005.09.011

Juston JM, DeBusk TA (2011) Evidence and implications of the background phosphorus concentration of submerged aquatic vegetation wetlands in Stormwater Treatment Areas for Everglades restoration. Water Resources Research 47:. https://doi.org/10.1029/2010WR009294

Juston JM, Kadlec RH (2019) Data-driven modeling of phosphorus (P) dynamics in low-P stormwater wetlands. Environmental Modelling & Software 118:226–240. https://doi.org/10.1016/j.envsoft.2019.05.002

Kadlec RH, Wallace SD (2009) Treatment wetlands. CRC Press, Boca Raton, FL

Liao C-Hsiang, Gurol MD (1995) Chemical Oxidation by Photolytic Decomposition of Hydrogen Peroxide. Environ Sci Technol 29:3007–3014. https://doi.org/10.1021/es00012a018

McCormick PV, Iii RBES, Backus JG, Kennedy WC (1998) Spatial and seasonal patterns of periphyton biomass and productivity in the northern Everglades, Florida, U.S.A. Hydrobiologia 362:185–210. https://doi.org/10.1023/A:1003146920533

McCormick PV, Laing JA (2003) Effects of increased phosphorus loading on dissolved oxygen in a subtropical wetland, the Florida Everglades. Wetlands Ecology and Management 11:199–216

McKenna JE (2003) Community metabolism during early development of a restored wetland. Wetlands 23:35–50. https://doi.org/10.1672/0277-5212(2003)023[0035:CMDEDO]2.0.CO;2

Mosisch TD, Bunn SE (1997) Temporal patterns of rainforest stream epilithic algae in relation to flow-related disturbance. Aquatic Botany 58:181–193. https://doi.org/10.1016/S0304-3770(97)00001-6

Newbold JD (1987) Phosphorus Spiralling in Rivers and River-Reservoir Systems: Implications of a Model. In: Craig JF, Kemper JB (eds) Regulated Streams. Springer US, pp 303–327

Newman S, Osborne TZ, Hagerthey SE, et al (2017) Drivers of landscape evolution: multiple regimes and their influence on carbon sequestration in a sub-tropical peatland. Ecol Monogr 87:578–599. https://doi.org/10.1002/ecm.1269

Newman S, Pietro K (2001) Phosphorus storage and release in response to flooding: implications for Everglades stormwater treatment areas. Ecological Engineering 18:23–38. https://doi.org/10.1016/S0925-8574(01)00063-5

NOAA (2013) CDMO NERR SWMP Data Management Manual. Belle W. Brauch Institute for Marine and Coastal Sciences, Georgetown, SC

Noe GB, Childers DL, Jones RD (2001) Phosphorus Biogeochemistry and the Impact of Phosphorus Enrichment: Why Is the Everglades so Unique? Ecosystems 4:603–624. https://doi.org/10.1007/s10021-001-0032-1

O’Donnell B, Hotchkiss ER (2019) Coupling Concentration- and Process-Discharge Relationships Integrates Water Chemistry and Metabolism in Streams. Water Resources Research. https://doi.org/10.1029/2019WR025025

Odum HT (1956) Primary production in flowing waters. Limnol Oceanogr 1:102–117

Osborne TZ, Bruland GL, Newman S, et al (2011) Spatial distributions and eco-partitioning of soil biogeochemical properties in the Everglades National Park. Environmental Monitoring and Assessment 183:395–408. https://doi.org/10.1007/s10661-011-1928-7

Pietro K (2012) Synopsis of the Everglades Stormwater Treatment Areas, Water Year 1996–2012. South Florida Water Management District, West Palm Beach, FL

Reddy KR, DeLaune RD (2008) Biogeochemistry of wetlands: science and applications. CRC Press, Boca Raton, FL

Reddy KR, Newman S, Osborne TZ, et al (2011) Phosphorous Cycling in the Greater Everglades Ecosystem: Legacy Phosphorous Implications for Management and Restoration. Critical Reviews in Environmental Science and Technology 41:149–186. https://doi.org/10.1080/10643389.2010.530932

Sand-Jensen K, Frost-Christensen H (1998) Photosynthesis of amphibious and obligately submerged plants in CO2-rich lowland streams. Oecologia 117:31–39. https://doi.org/10.1007/s004420050628

Sand-Jensen K, Prahl C, Stokholm H (1982) Oxygen Release from Roots of Submerged Aquatic Macrophytes. Oikos 38:349–354. https://doi.org/10.2307/3544675

South Florida Water Management District (2013) Restoration Strategies Regional Water Quality Plan: Science Plan for the Everglades Stormwater Treatment Areas. South Florida Water Management District, West Palm Beach, FL

Staehr PA, Sand-Jensen K (2007) Temporal dynamics and regulation of lake metabolism. Limnol Oceanogr 52:108–120. https://doi.org/10.4319/lo.2007.52.1.0108

Staehr PA, Sand-Jensen K, Raun AL, et al (2010) Drivers of metabolism and net heterotrophy in contrasting lakes. Limnol Oceanogr 55:817–830. https://doi.org/10.4319/lo.2010.55.2.0817

Staehr PA, Testa JM, Kemp WM, et al (2012) The metabolism of aquatic ecosystems: history, applications, and future challenges. Aquatic Sciences 74:15–29. https://doi.org/10.1007/s00027-011-0199-2

Stanley EH, Johnson MD, Ward AK (2003) Evaluating the influence of macrophytes on algal and bacterial production in multiple habitats of a freshwater wetland. Limnology and Oceanography 48:1101–1111. https://doi.org/10.4319/lo.2003.48.3.1101

Stevenson RJ, Bothwell ML, Lowe RL (eds) (1996) Algal Ecology: Freshwater Benthic Ecosystem. Academic Press

Thébault J, Schraga TS, Cloern JE, Dunlavey EG (2008) Primary production and carrying capacity of former salt ponds after reconnection to San Francisco Bay. Wetlands 28:841–851. https://doi.org/10.1672/07-190.1

Tsai J-W, Kratz TK, Hanson PC, et al (2011) Metabolic changes and the resistance and resilience of a subtropical heterotrophic lake to typhoon disturbance. Canadian Journal of Fisheries and Aquatic Sciences 68:768–780. https://doi.org/10.1139/f2011-024

Tuttle CL, Zhang L, Mitsch WJ (2008) Aquatic metabolism as an indicator of the ecological effects of hydrologic pulsing in flow-through wetlands. Ecological Indicators 8:795–806. https://doi.org/10.1016/j.ecolind.2007.09.005

UF-WBL (2017) Evaluation of Soil Biogeochemical Properties Influencing Phosphorus Flux in the Everglades Stormwater Treatment areas: 2016-2017 Annual Report. University of Florida, Gainesville, FL

Updegraff K, Pastor J, Bridgham SD, Johnston CA (1995) Environmental and Substrate Controls over Carbon and Nitrogen Mineralization in Northern Wetlands. Ecological Applications 5:151–163. https://doi.org/10.2307/1942060

Vannote RL, Minshall GW, Cummins KW, et al (1980) The River Continuum Concept. Can J Fish Aquat Sci 37:130–137. https://doi.org/10.1139/f80-017

Villapando O, King J (2018) Appendix 5C-3: Evaluation of Phosphorus Sources, Forms, Flux, and Transformation Processes in the Stormwater Treatment Areas. In: 2018 South Florida Environmental Report. South Florida Water Management District, West Palm Beach, FL

Webster JR (2007) Spiraling down the river continuum: stream ecology and the U-shaped curve. Journal of the North American Benthological Society 26:375–389. https://doi.org/10.1899/06-095.1

White JR, Reddy KR, Majer-Newman J (2006) Hydrologic and Vegetation Effects on Water Column Phosphorus in Wetland Mesocosms. Soil Science Society of America Journal 70:1242. https://doi.org/10.2136/sssaj2003.0339

Yarwood SA (2018) The role of wetland microorganisms in plant-litter decomposition and soil organic matter formation: a critical review. FEMS Microbiology Ecology 94:. https://doi.org/10.1093/femsec/fiy175

Zhao H, Piccone T (2018) Appendix 5C-6: Summary Report for Stormwater Treatment Area 2 Flow-ways 1, 2, and 3 Water and Total Phosphorus Budget Analyses. In: South Florida Environmental Report, 2018th edn. South Florida Water Management District, West Palm Beach, FL

