## Supplementary material for "Understanding potential drivers of aquatic metabolism in a subtropical treatment wetland": Supplmental

Table S1. Summary of parameters, matrices, analytical methods and minimum detection limit (MDL) used for this study. Additional parameters were collected as part of the larger study (Reddy 2017) but not used in this study. All analytical methods are consistent with Florida Department of Environmental Protection or U.S. Environmental Protection Agency Standard Operating Procedures and methods.

| **Matrix** | **Parameter** | **Abbreviation** | **Analytical Method** | **Minimum Detection Limit** | **Method** |
| --- | --- | --- | --- | --- | --- |
| Surface Water | Total Phosphorus | TP | SM4500PF | 2 µg P L^-1^ | Clesceri et al. (1998) |
|  | Chlorophyll-*a* | TN | SM4500NC | 0.18 µg L^-1^ | Arar (1997) |
|  | Dissolved Organic Carbon | DOC | SM5310B | 0.8 mg C L^-1^ | Clesceri et al. (1998) |
| Flocculent and Soil | Loss-on-ignition^1^ | LOI | Calculation^2^ | 1.0 % | **---** |
|  | Total Carbon | TC | SFWMD 3200 | 2 g C kg^-1^ | SFWMD (2015) |

^1^ Loss-on-ignition was calculated from the difference between 100% and percent ash determined by the analytical method identified as SFWMD 1610 (SFWMD 2015)**.**

^2^ Table References

Arar EJ (1997) Method 447.0 - Determination of Chlorophylls a and b and Identification of Other Pigments of Interest in Marine and Freshwater Algae Using High Performance Liquid Chromatography with Visible Wavelength Detection. United States Environmental Protection Agency, Washington DC

Clesceri LS, Greenberg AE, Eaton AD (eds) (1998) Standard Methods for the Examination of Water and Wastewater. American Public Health Association

Reddy K (2017) Evaluation of Soil Biogeochemical Properties Influencing Phosphorus Flux in the Everglades Stormwater Treatment areas: 2016-2017 Annual Report. University of Florida, Gainesville, FL

SFWMD (2015) Chemistry Laboratory Quality Manual. South Florida Water Management District, West Palm Beach, FL

Table S2**.** Event characteristics including duration, hydraulic and phosphorus loading rates (HLR and PLR, respectively) and median detection time of the five flow events for STA-2 flow-ways (FWs) 1 and 3.

| **STA Flow-way** | **Event** | **Start - End Date** | **Duration (Days)** | **HLR**  **(cm d⁻¹)** | **PLR**  **(mg m⁻² d⁻¹)** | **Median**  **Detention**  **Time (d)** |
| --- | --- | --- | --- | --- | --- | --- |
| FW 1 | 1 | Aug 10 - Sep 14, 2015 | 35 | 0.56 ± 0.12 | 0.46 ± 0.10 | 35.7 |
| FW 1 | 2 | Oct 20 - Nov 29, 2015 | 40 | 1.20 ± 0.25 | 0.37 ± 0.08 | 27.1 |
| FW 3 | 3 | Feb 22 - Apr 11, 2016 | 49 | 3.17 ± 0.63 | 1.45 ± 0.30 | 2.3 |
| FW 3 | 4 | Jun 27 - Aug 29, 2016 | 63 | 2.00 ± 0.24 | 1.16 ± 0.16 | 16.0 |
| FW 3 | 5 | Oct 12 - Nov 22, 2016 | 41 | 5.88 ± 0.84 | 4.33 ± 0.68 | 5.2 |
| FW 1 | 6 | May 29 - Jul 31, 2017 | 63 | 3.84 ± 0.71 | 7.10 ± 1.39 | 25.0 |

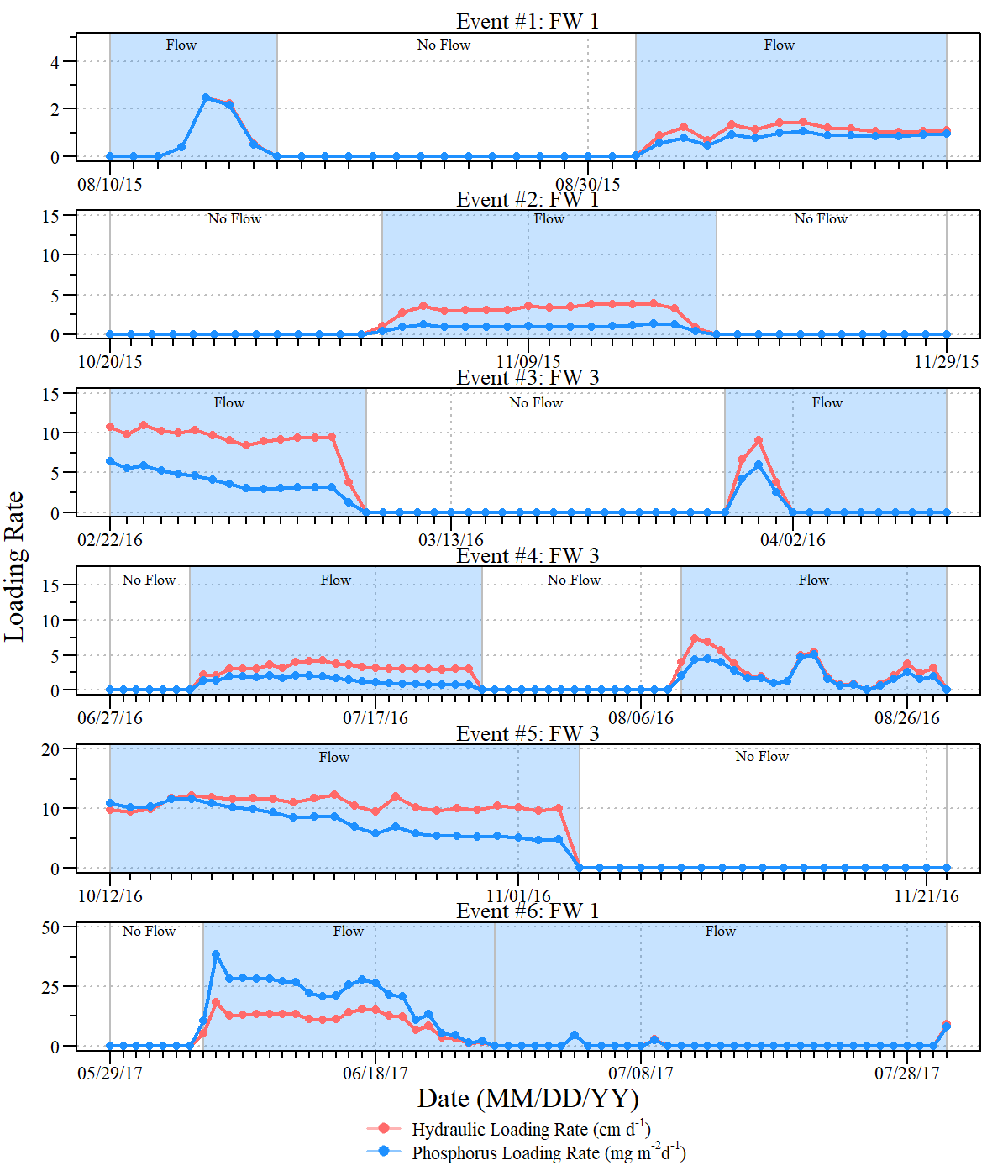

Figure S1. Hydrologic and Phosphorus loading rates (HLR and PLR, respectively) for the six flow events within flow-ways (FWs) 1 and 3 of Stormwater Treatment Area-2 between August 10^th^, 2015 and July 31^st^, 2017.

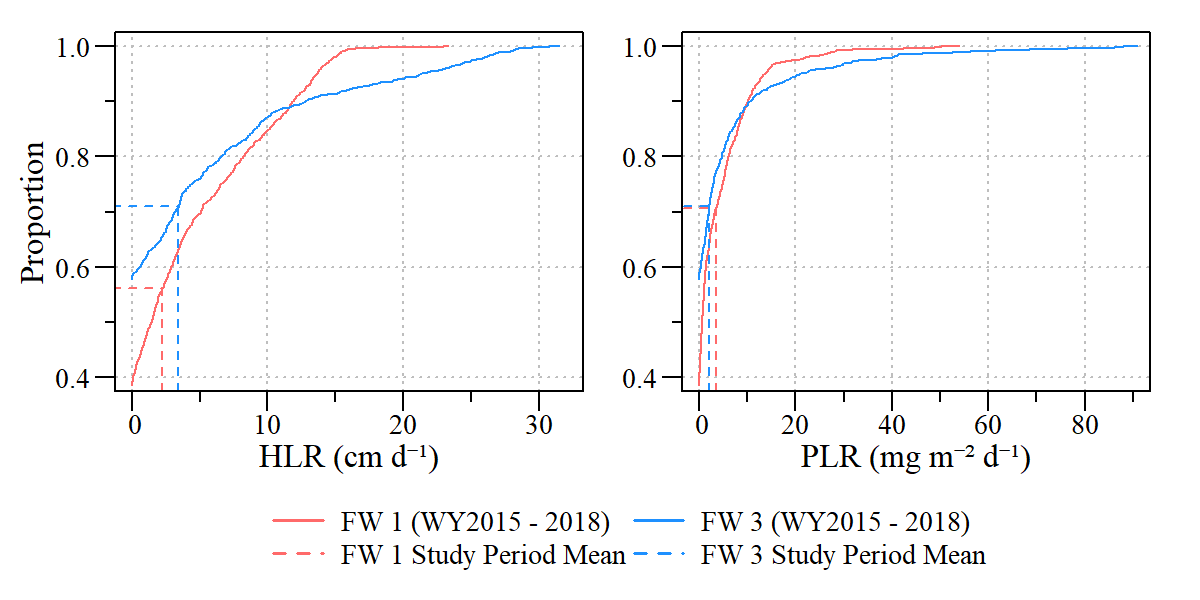

Figure S2. Cumulative distribution plots of hydraulic (left) and phosphorus (right) loading rates for flow-ways 1 and 2 (FW1 and FW2, respectively) for data collected between May 1^st^ 2014 and April 30^th^ 2018 (solid lines) relative to experiment flow period mean values (dashed values) for each flow-way.

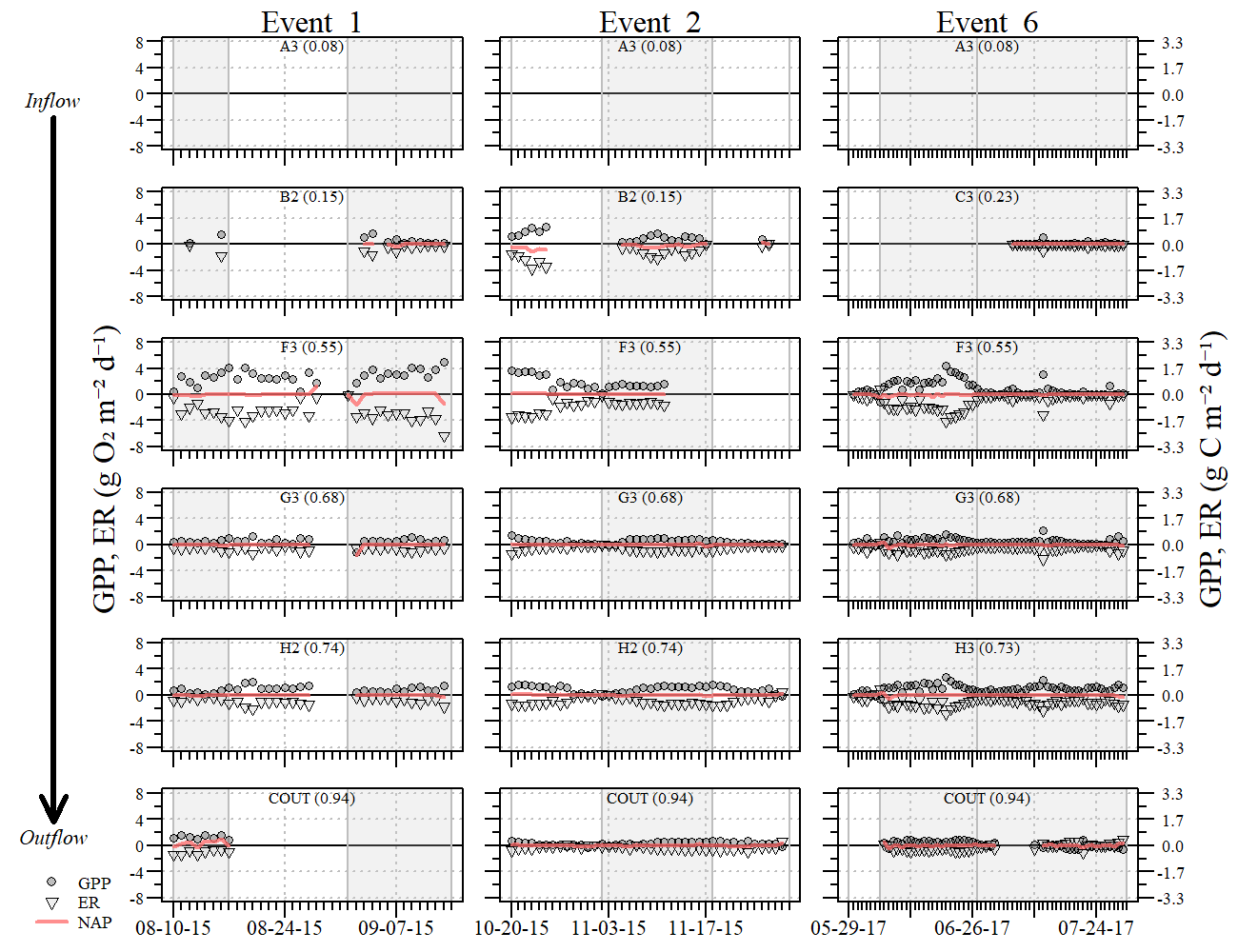

Figure S3. Daily gross primary productivity (GPP; circles), ecosystem respiration (ER; inverted triangle) and net aquatic productivity (NAP; red-line) along the Stormwater Treatment Area 2 flow-way 1 transect, moving from inflow to outflow, each plot is identified with site names and fractional distance downstream (0.0 = inflow; 1.0 = outflow) during three flow events.

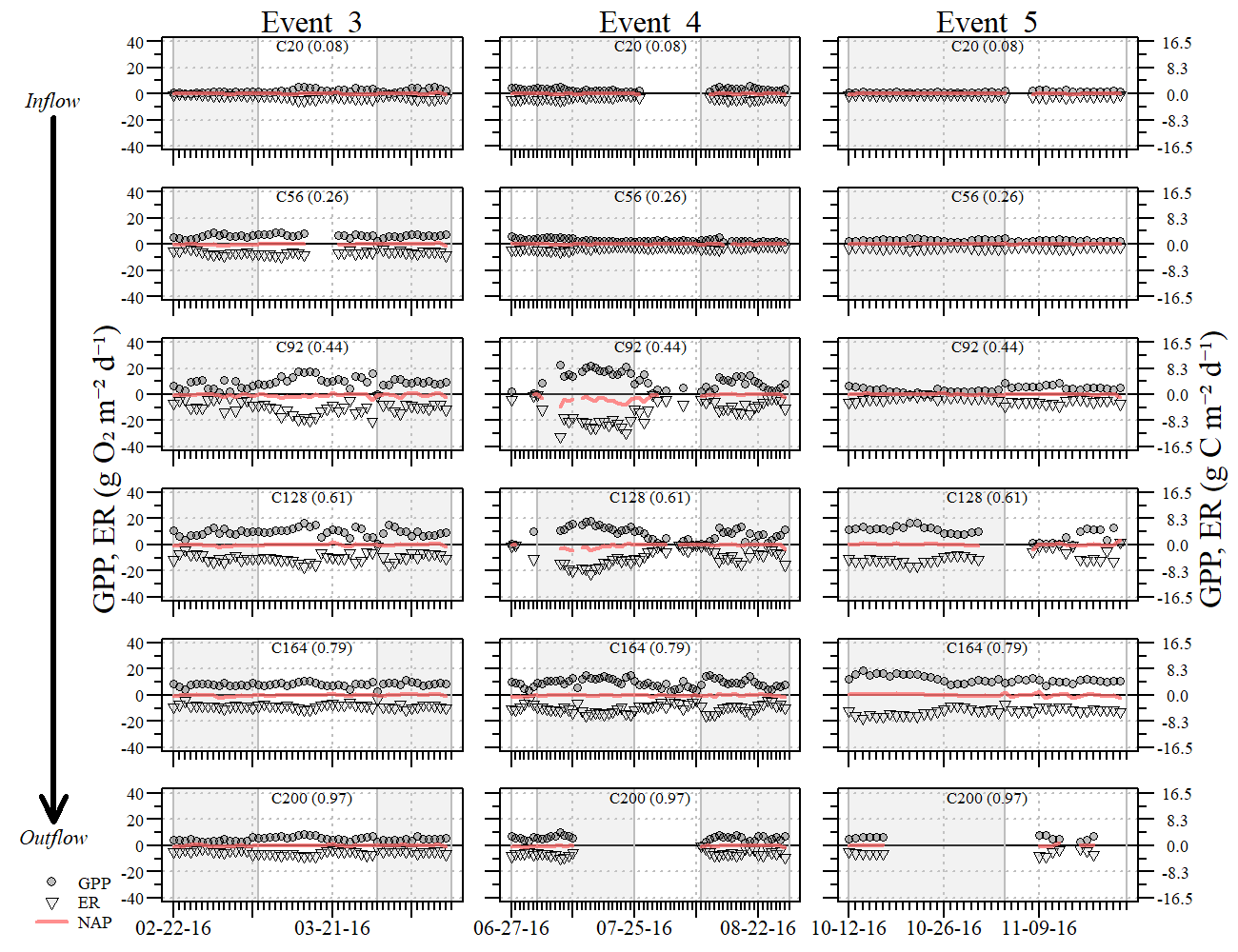

Figure S4. Daily gross primary productivity (GPP; circles), ecosystem respiration (ER; inverted triangle) and net aquatic productivity (NAP; red-line) along the Stormwater Treatment Area 2 flow-way 3 transect, moving from inflow to outflow, each plot is identified with site names and fractional distance downstream (0.0 = inflow; 1.0 = outflow) during three flow events.

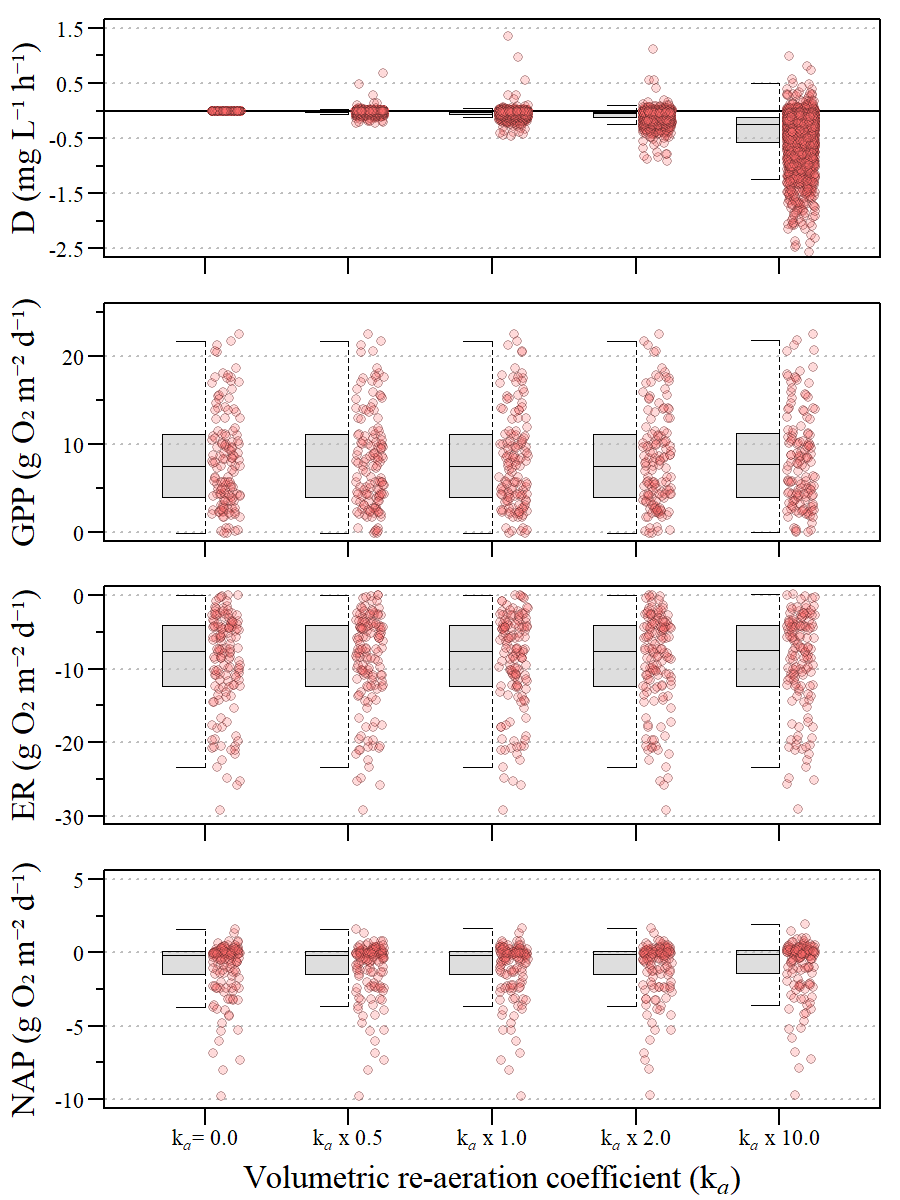

Figure S5. Sensitivity analysis of the volumetric re-aeration coefficient used to compute diffusive oxygen uptake (D; Eq 2), gross primary production (GPP), ecosystem respiration (ER) and net aquatic productivity (NAP). This data is specific to site C92 in flow-way 3 as this site had the most data across the study period and the greatest variability in metabolism metrics (Fig S3 and S4).
